# Early-stage cancer results in a multiplicative increase in cell-free DNA originating from healthy tissue

**DOI:** 10.1101/2024.01.26.577500

**Authors:** Konstantinos Mamis, Ivana Bozic

**Author notes:** Corresponding author (IB).

## Abstract

Cell-free DNA is a promising biomarker for cancer detection. However, the sources of elevated cell-free DNA (cfDNA) in patients with early-stage cancer, and the mechanisms by which cfDNA is shed into, and subsequently cleared from the circulation are still poorly understood. Leveraging a rich dataset of cfDNA in healthy individuals and early-stage cancer patients, we find a multiplicative increase in cfDNA concentration in the presence of cancer. This increase is cancer type-specific, ranging from a ∼1.3-fold increase in lung cancer, to a ∼12-fold increase in liver cancer, and does not originate from tumor, but from healthy tissue. Employing an additional dataset reporting the tissue of origin of cfDNA, we observe a significant increase in the correlation between cfDNA originating from leukocytes and from non-leukocyte sources in cancer patients. Introducing a mathematical model for cfDNA dynamics, we find that the observed correlation can be explained by a saturation mechanism in cfDNA clearance. Saturation in clearance implies that smaller increases in cfDNA shedding may lead to proportionally larger increases in cfDNA levels. Our findings quantify cfDNA dynamics in patients with cancer, with implications for improving the accuracy of liquid biopsies for early cancer detection.

## Introduction

Successful detection of cancer in early stages results in more effective treatment, and significantly improved survival prognosis. Identifying biomarkers that can reliably indicate the presence of early-stage cancers is one of the major challenges in oncology [1]. One such promising biomarker is cell-free DNA (cfDNA), composed of the DNA fragments not bound to cells, that can be detected in the circulation. The presence of cfDNA in the blood was first discovered by Mandel and Metais [2] in the 40’s; since then, increased levels of cfDNA concentrations have been reported in cancer patients [3–5], especially in advanced and metastatic stages [6]. Furthermore, Stroun *et al*. [7] uncovered in the 80’s that cfDNA from cancer patients harbored mutations that were also present in patients’ tumors.

These discoveries motivated further research on detecting cancer by performing “liquid biopsies” on cfDNA, as a less intrusive alternative to the biopsy on the tumor itself. However, the mechanisms by which DNA is shed into, and subsequently cleared from the circulation are not fully understood [8–10].

The cfDNA clearance occurs primarily in the liver (70%-90%), mediated by Kupffer cells, with additional contributions from kidneys and the spleen [11]. Clearance is rapid, with reported cfDNA half-life estimates ranging from approximately 15 minutes to 2.5 hours [12–15]. In settings of impaired cfDNA clearance, such as sepsis [16] or, in the cancer context, the pharmacologic attenuation of cfDNA clearance in tumor-bearing mice [17], cfDNA levels rise without a significant change in composition (tissue-of-origin in sepsis; tumor fraction in mouse experiments), indicating that clearance acts non-selectively on cfDNA. Importantly, clearance capacity is finite and becomes saturated at high concentrations, as demonstrated by dose-dependent hepatic uptake of radiolabeled mononucleosomes in mice [18]. These mechanisms are critical for interpreting cfDNA dynamics under cancer conditions, yet their quantitative impact remains incompletely characterized.

Recent work demonstrates that the majority of cfDNA in cancer patients with highly elevated cfDNA originates from leukocytes, and that the presence of cancer likely results in a systemic increase in cfDNA concentration in these patients [19]. Whether a similar systemic increase due to cancer is present in all cancer patients is currently not known, as many patients, in particular those with early-stage cancer, do not exhibit elevated cfDNA levels.

In this setting, several questions must be answered in order to better understand the mechanics of DNA circulating in plasma, which could lead to improving the accuracy of liquid biopsy screening tests. In what way, and how much does cancer presence affect the total cfDNA concentration? Is this effect cancer type-specific, and does it differ between early cancer stages? In the present study, we address these questions through a novel analysis of a large cfDNA dataset from Cohen *et al*. [20], and quantify the differences in cfDNA dynamics between healthy individuals and patients with stage I-III ovarian, liver, stomach, pancreatic, esophageal, colorectal, lung, and breast cancer. Employing an additional dataset from Mattox *et al*. [19] reporting the tissue of origin for cfDNA in healthy individuals and pancreatic and ovarian cancer patients, we are able to provide insight regarding the mechanisms by which cfDNA is cleared from circulation, and estimate the increases in cfDNA shedding from leukocytes and non-leukocyte sources due to cancer.

## Results

Cohen et al. [20] reported cfDNA concentrations in plasma for healthy individuals and cancer patients, as well as the concentration and mutant allele frequency (MAF) of DNA fragments in plasma harboring mutations in cancer driver genes; we refer to this portion of cfDNA as circulating mutant DNA (cmDNA) (Materials and Methods, Table S1). Among the healthy individuals included in our analysis, cmDNA was detected in 90.6% of samples, with a median cfDNA concentration of 2.83 ng/mL, whereas samples without detectable mutations showed much lower cfDNA (median 0.05 ng/mL; S1 Fig.). As we aim to quantify and compare the effect of cancer on both cfDNA and cmDNA concentrations, we focus on the 90.6% of healthy individuals with cmDNA detected in plasma. Full exclusion criteria and cohort sizes for healthy individuals and cancer patients are provided in Materials and Methods.

### Multiplicative increase in cfDNA originating from healthy tissue

Patients with late-stage cancer typically have cfDNA concentrations that are significantly higher than those in healthy individuals [6,10]. However, cfDNA concentrations in patients with early-stage cancer often overlap with those found in healthy individuals (Fig. 1). Performing the Kolmogorov-Smirnov (K-S) statistical test to determine the similarity of probability distributions of cfDNA concentration in healthy individuals and cancer patients, we find that the presence of stage I-III lung, breast, colorectal, pancreatic, ovarian, esophageal, stomach and liver cancer leads to a multiplicative increase in cfDNA concentration compared to healthy individuals (Fig. 2). In other words, we find that the entire dataset of cfDNA concentrations is multiplied by a positive factor *α* in patients with cancer compared to healthy individuals (Kolmogorov-Smirnov test at 5% significance level, Materials and Methods). This effect can be seen as a horizontal “shift” or translation when the cumulative distribution function (CDF) is plotted on a logarithmic *x*-scale (Fig. 2).

**Figure 1.**
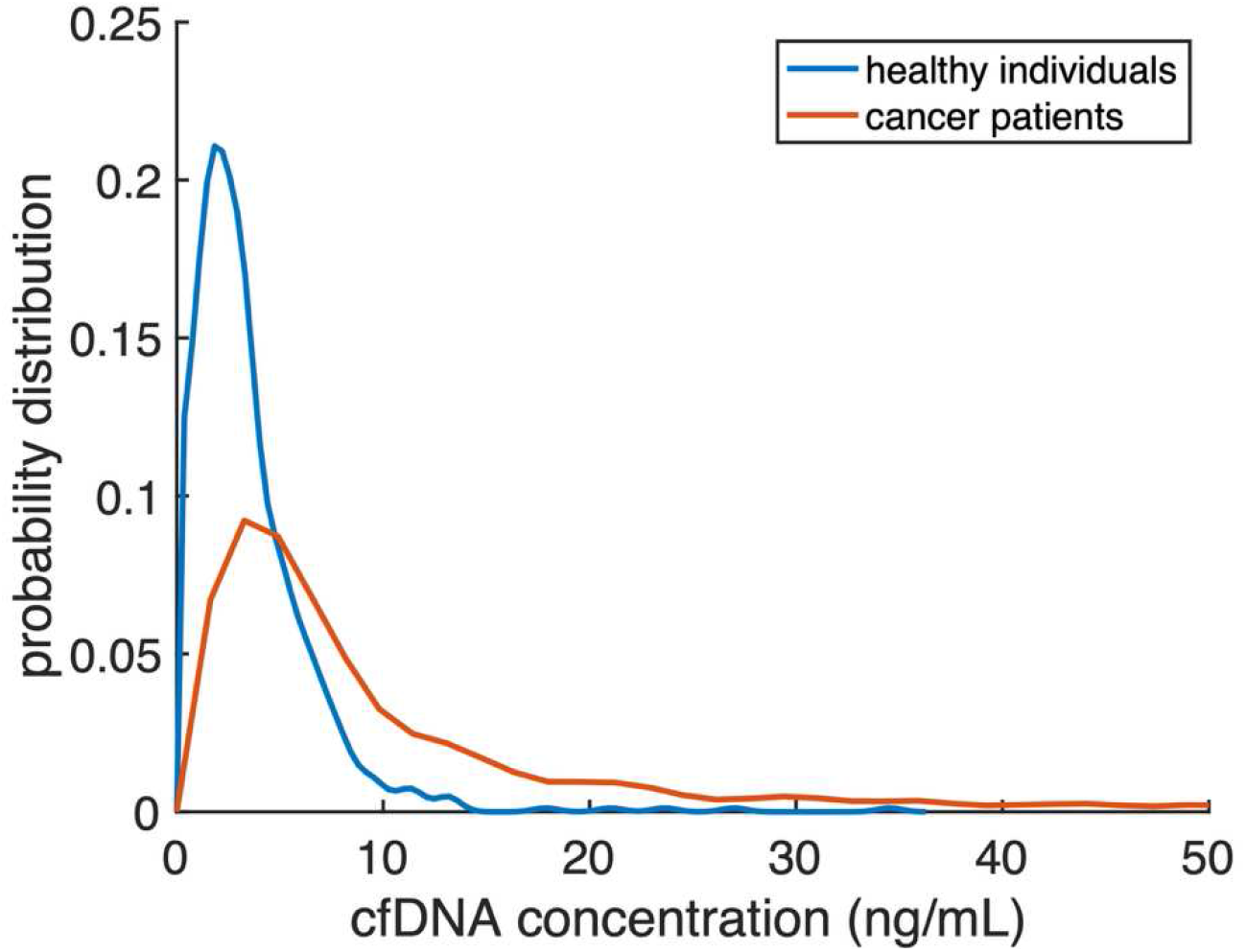
Probability distribution of cfDNA concentrations in healthy individuals and patients with stage I-III cancer. Probability density functions for the cfDNA concentration in healthy individuals with mutations detected in plasma (*N*=572) and patients with lung, breast, colorectal, pancreatic, ovarian, esophageal, stomach and liver cancer in stages I-III (*N*=952), as reported in Cohen *et al*. (2018) [20].

**Figure 2.**
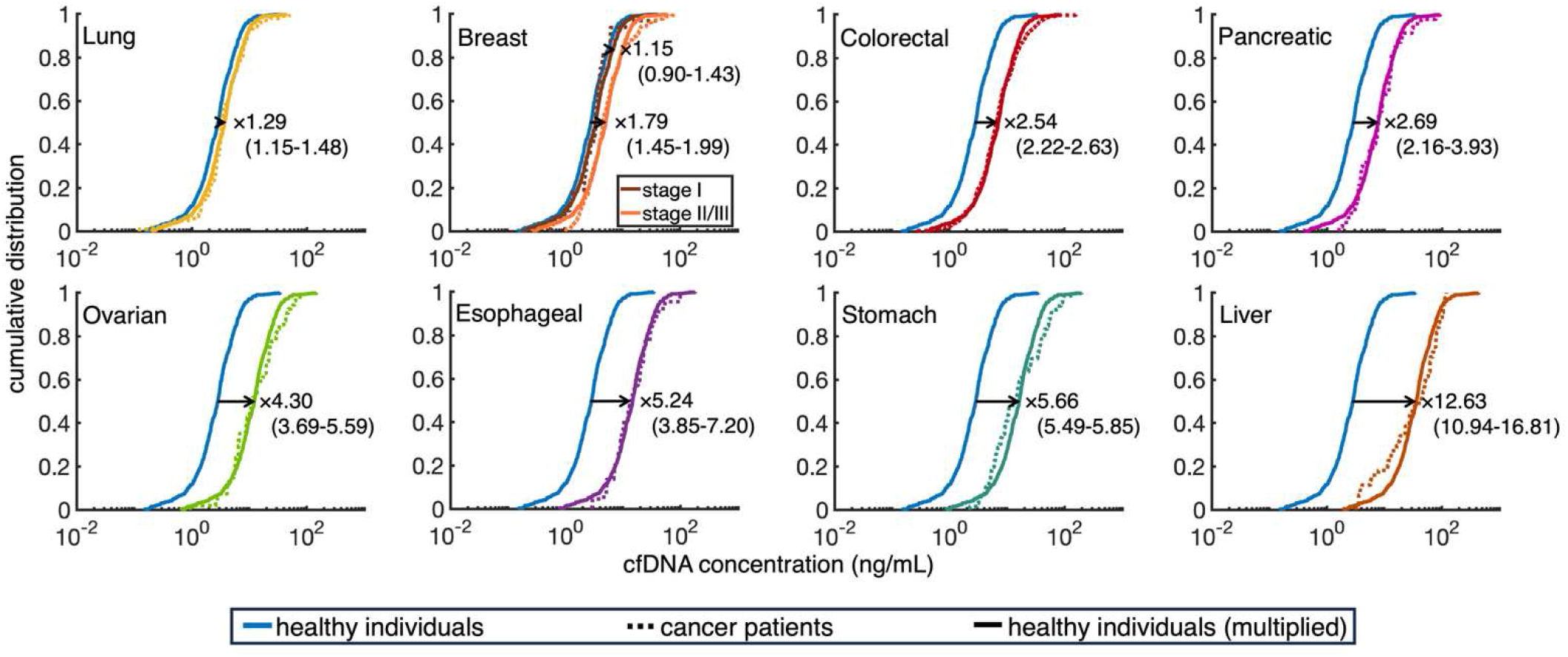
Quantifying the multiplicative increase *α* in cfDNA concentration in the presence of cancer. Cumulative distribution functions for the cfDNA concentration in healthy individuals with mutations detected in plasma (*N*=572) and patients with lung (*N*=103), stage I breast (*N*=32), stage II/III breast (*N*=177), colorectal (*N*=385), pancreatic (*N*=47), ovarian (*N*=54), esophageal (*N*=44), stomach (*N*=66) and liver (*N*=44) cancer. The value and range shown on each panel represent the multiplicative increase *α* and its range of values not rejected by the K-S test at the 5% significance level. Median empirical cfDNA concentrations (corresponding to 0.5 value of cumulative distribution) are 2.83 ng/mL for healthy individuals, and 3.31, 3.42, 4.60, 6.62, 7.76, 11.18, 13.97, 14.14, and 42.27 ng/mL for patients with lung, stage I breast, stage II/III breast, colorectal, pancreatic, ovarian, esophageal, stomach and liver cancer, respectively.

The multiplicative increase *α* is specific to each cancer type: 1.29 (value range not rejected by K-S test at 5% significance level: 1.15-1.48) for lung cancer, 1.15 (0.9-1.43) for stage I breast cancer, 1.79 (1.45-1.99) for stage II/III breast cancer, 2.54 (2.22-2.63) for colorectal, 2.69 (2.16-3.93) for pancreatic, 4.3 (3.69-5.59) for ovarian, 5.24 (3.85-7.2) for esophageal, 5.66 (5.49-5.85) for stomach, and 12.63 (10.94-16.81) for liver cancer (Fig. 1, S2 Fig., S2 Table, Materials and Methods). Additionally, as the visual counterpart of the K-S test, we construct the Probability–Probability (P–P) plots, which compare the CDFs of two datasets (Materials and Methods). For each cancer type, the P–P plots show strong concordance between the empirical CDFs for cfDNA concentrations in cancer patients and in healthy individuals multiplied by *α* (Fig. S2). P-P plots for stomach and liver cancer diverge slightly from the diagonal due to greater differences at the distribution tails between cancer and scaled healthy CDFs (Fig. 2). P-P plots amplify tail effects, but the overall agreement supports the validity of the shift.

There are no statistically significant differences in cfDNA concentration between stages I to III, for all cancer types except for breast and liver cancer; however, the data for stage I pancreatic, esophageal and liver cancer is limited (2, 5 and 5 patients respectively). We found no significant differences in cfDNA concentrations between sexes. Although the median age of healthy individuals was younger than that of cancer patients (36.5 years compared to 57–68 years; Table S1b), there was no statistical difference in cfDNA levels between the 288 healthy individuals under 40 years and the 263 healthy individuals over 50 years of age (K-S test p > 0.05, S3 Figure). We selected the < 40 y and > 50 y age groups of healthy individuals to reflect the previously reported median ages (36.5 y in healthy individuals and 57-68 y in cancer patients) and to test whether an age gap of that size is associated with significantly different cfDNA levels within healthy individuals. The intermediate 40-50 y group is small (n = 21) and was not analyzed separately. Furthermore, we grouped all healthy individuals into ≤ 40 y and > 40 y age groups; again, the Kolmogorov-Smirnov test showed no significant differences in cfDNA levels between these two age groups.

To determine if a multiplicative increase in cfDNA concentration is also present in the earliest cancer cases, we calculate *α* between cfDNA concentrations in healthy individuals and cancer patients, restricting the cancer data to only stage I cases. We do not perform the calculation for pancreatic, esophageal and liver cancer, due to insufficient number of patients in stage I. Breast cancer patients have already been analyzed separately in stage I and stage II-III due to the significant difference in cfDNA concentrations between these groups. We find that *α* is 1.33 (1.04-1.71) for lung cancer, 2.39 (2.02-2.88) for colorectal cancer, 6.99 (3.64-14.82) for ovarian cancer, and 4.83 (2.27-6.38) for stomach cancer (S4 Fig.). These ranges for *α* overlap with the ranges calculated when all stage I-III cancer patients were considered. We note that, for ovarian cancer, the multiplicative factor is 6.99 (3.64–14.82) for stage I compared to 4.30 (3.69–5.59) for stages I–III. The apparently higher factor for stage I ovarian cancer reflects the small sample size (N = 9), where a few patients with elevated cfDNA disproportionately influence the estimate, resulting in a substantially wider range. Importantly, the ranges for stage I overlap with those for stages I–III, demonstrating consistency of the multiplicative factor between stage I and all stages combined. Additionally, cfDNA concentrations do not differ significantly between stage I and all stages combined in all four cancer types considered.

We next asked if the source of the increased cfDNA concentration in early-stage cancer patients is the cancer itself or if the increase is originating from the healthy tissue. To address this, we analyzed 484 patients from the Cohen et al. [20] dataset whose tumors harbored at least one high-frequency mutation (MAF ≥30%). MAF in plasma reflects both the clonality of the mutation in its tissue of origin and that tissue’s contribution to total cfDNA. For example, if a mutation has a MAF in the tumor of 30%, and if 50% of cfDNA originates from the tumor, then, on average, the mutation should have MAF around 15% in plasma. Even if the biopsy overrepresents the mutation compared to the entire tumor -for example, MAF of 30% in the biopsy versus 10% in the tumor-if 50% of cfDNA originates from the tumor, the expected plasma MAF would still be around 5%. The exact numbers may be different in some cases due to differences in copy number between cancer and healthy tissue, but are expected to exhibit similar behavior. We recognize that clearance dynamics could influence absolute cfDNA levels; however, experimental evidence indicates that impaired clearance increases cfDNA globally without altering its composition [16,17], suggesting that clearance differences do not systematically bias the relative contribution of tumor versus healthy tissue to plasma cfDNA. Therefore, if most of cfDNA originates from the tumor, the high-MAF mutations in the tumor should appear in plasma with significant MAF (≫1%). However, in these 484 patients, only 3.1% of top plasma mutations in cancer patients have MAF above 10%, 5.8% above 5%, and 13.4% above 1% (S5 Fig., red line). These findings indicate that, even when tumors carry high-clonality mutations, most cfDNA in plasma does not originate from the tumor but from non-tumor (healthy) tissue.

The expectation that a mutation with high MAF in the tumor should be present in plasma with significant MAF (≫1%), assuming that most of cfDNA originates from the tumor, holds even if the mutation was missed by the tumor biopsy, provided it is theoretically detectable in plasma (i.e. included in the mutational panel). For that reason, we also examined the distribution of plasma mutation MAFs for the 162 cancer patients whose tumors harbored only mutations with MAF < 30% (S5 Fig., blue line). The results were similar to the distribution of plasma MAFs for patients with tumor MAF≥30%; in fact, the two groups were indistinguishable, as confirmed by a Kolmogorov–Smirnov test (p > 0.05).

### Multiplicative increase in non-tumor circulating mutant DNA

We next hypothesized that, since cfDNA is mainly shed from non-tumor sources and exhibits a multiplicative increase in the presence of cancer, then a similar increase should also be observed in circulating mutant DNA (cmDNA) that does not originate from tumor. In the dataset reported by Cohen et al. [20], which includes a total of 959 cancer patients with stage I–III disease across eight cancer types, information on both the top mutation detected in plasma and the mutations identified in tumor is available for 645 patients (Materials and Methods, S1c Table). The cmDNA concentration reported by Cohen et al. [20] corresponds to the top plasma mutation. When the top mutation detected in plasma is not found among the mutations reported in the tumor biopsy, we conclude that this mutation most likely does not originate from the tumor, but rather from healthy tissue. Comparing the cmDNA concentration in healthy individuals and in cancer patients in which cmDNA detected in plasma is not detected in tumor (S1c Table), we find increases by a multiplicative factor for every cancer type: 1.92 (1.42-2.52) for lung, 2.42 (2.12-2.76) for breast, 2.36 (1.92-2.66) for colorectal, 4.01 (1.55-7.34) for pancreatic, 3.31 (1.76-5.38) for esophageal, 4.38 (2.74-5.77) for stomach, and 12.37 (5.54-24.62) for liver cancer (Fig. 3, S6 Fig., S3 Table, Materials and Methods). We do not calculate the multiplicative factor for ovarian cancer, since the dataset reported only one ovarian cancer patient with cmDNA not originating from the tumor.

**Figure 3.**
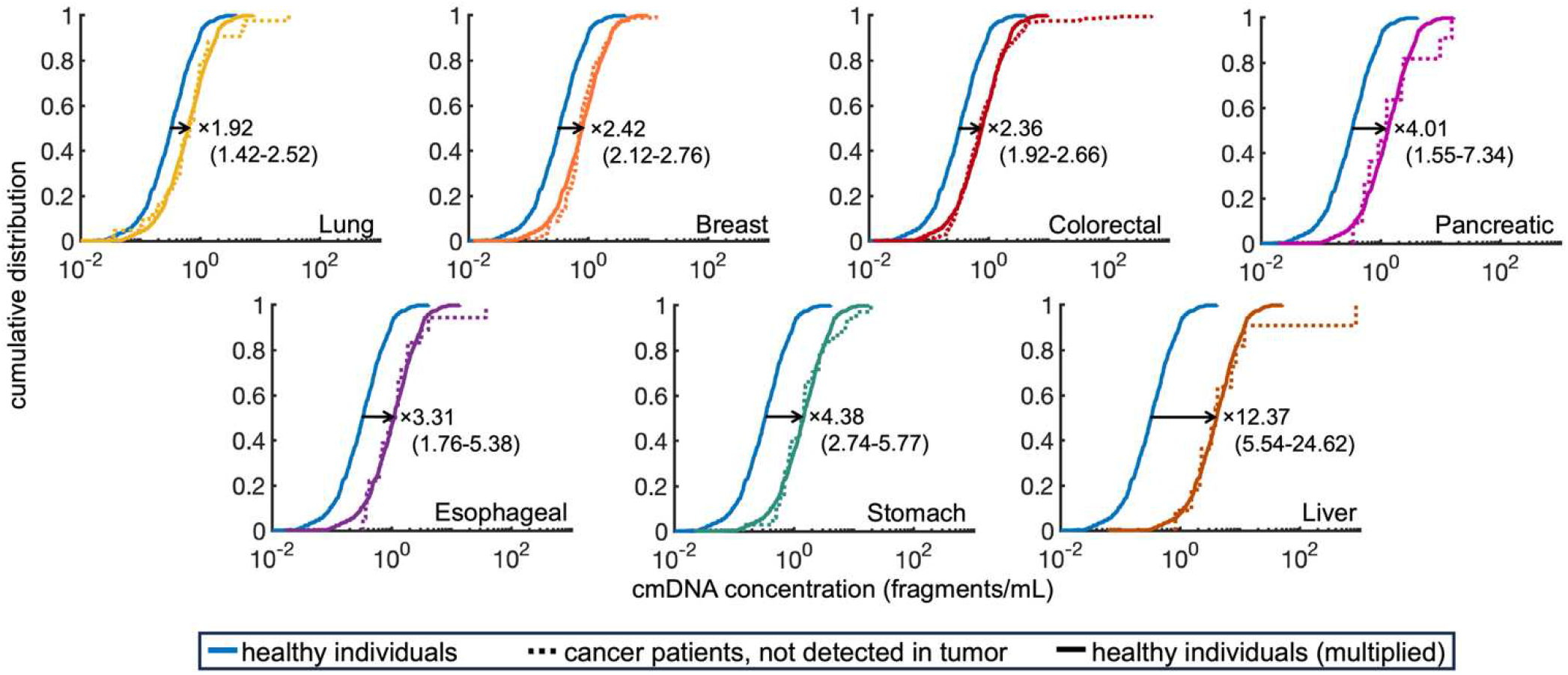
Mutations detected in plasma of cancer patients but not detected in tumor exhibit a multiplicative increase in concentration compared to mutations detected in plasma of healthy individuals. Cumulative distribution functions for the concentration of cmDNA found in plasma of healthy individuals (*N*=572) and patients with lung (*N*=43), breast (*N*=90), colorectal (*N*=205), pancreatic (*N*=11), esophageal (*N*=18), stomach (*N*=34) and liver (*N*=11) cancer in which the mutation in plasma was not detected in tumor. The value and range shown on each panel represent the multiplicative increase *α* and its range of values not rejected by the K-S test at the 5% significance level. Median empirical cmDNA concentrations (corresponding to 0.5 value of cumulative distribution) are 0.33 fragments/mL for healthy individuals, and 0.67, 0.73, 0.72, 1.23, 1.05, 1.39, and 3.85 fragments/mL for patients with lung, breast, colorectal, pancreatic, esophageal, stomach and liver cancer, respectively.

Mutant DNA in plasma may originate from healthy or tumor tissue; however, parallel sequencing polymerase chain reaction (PCR) methods may introduce artifactual mutations with low MAFs. Circulating mutant DNA data from Cohen *et al*. [20] which we analyze in this work were generated using the SafeSeqS approach for PCR, developed to reduce the introduction of artifactual mutations [21]. In subsequent work [22], Cohen *et al*. report that the artifactual mutations in plasma analyzed by SafeSeqS method appeared with a MAF lower than 0.013%. Hence, we perform the cmDNA analysis again, disregarding the mutations with low MAFs; the results remain similar to the results obtained for all mutations (S7 Fig., S3 Table, Materials and Methods).

### Insight into cfDNA dynamics

The multiplicative increase in cfDNA levels observed between healthy and cancer patients can be the result of increased cfDNA shedding from non-tumor sources, a slower rate of cfDNA clearance, or a combination of both. In order to provide insight regarding the cfDNA dynamics and the cause of cfDNA increase in early-stage cancer, we utilize data from Mattox *et al*. [19] regarding the cfDNA tissue of origin for 64 healthy individuals, 79 patients with stage I-III pancreatic cancer, and 32 patients with stage I-III ovarian cancer. The identification of cfDNA tissue of origin in Mattox *et al*. [19] was based on the deconvolution of the cfDNA methylation profile using two different reference matrices for the tissue-specific methylation regions, as determined by Sun *et al*. [23] and Moss *et al*. [24], and two different deconvolution algorithms (quadratic programming or non-linear least squares). This results in four different cfDNA compositions reported for each individual from this dataset.

First, for this dataset, we perform the same statistical analysis to determine the multiplicative factor between healthy and cancer patients, finding an *α* of 4.4 (3.8-4.81) and 2.5 (2.1-3.33) for pancreatic and ovarian cancer respectively. The ranges of multiplicative factor values as calculated from the datasets of Mattox *et al*. [19] and Cohen *et al*. [20] are overlapping for pancreatic cancer; for ovarian cancer they are comparable yet non-overlapping, with a lower multiplicative factor calculated from the dataset of Mattox *et al*. [19].

In healthy individuals from Mattox *et al*. [19], most cfDNA originates from leukocytes; the different deconvolution methods employed report a median 66.8%-78% of cfDNA from leukocytes. In pancreatic and ovarian cancer patients, the cfDNA percentage from leukocytes is similar (median 65.1%-70.8% and 70.3%-75.5% respectively), in cases where the reference deconvolution matrix by Sun *et al*. [23] is used. Use of the reference deconvolution matrix by Moss *et al*. [24] results in decreased percentages of cfDNA from leukocytes (∼53% in pancreatic cancer and 52%-58.5% in ovarian cancer patients).

From this dataset, we also calculate the Pearson correlation coefficient for cfDNA concentrations from leukocytes and non-leukocyte sources, after we exclude outliers (three outliers with high cfDNA concentrations from healthy individuals and one from ovarian cancer patients, S1 Text). Note that the exclusion of outliers does not have a significant effect on the multiplicative factors calculated above. Correlation coefficient in healthy individuals is calculated from -0.29 to 0.20, depending on the cfDNA deconvolution method, 0.31-0.58 in pancreatic cancer patients, and 0.32-0.87 in ovarian cancer patients. Thus, apart from the multiplicative increase in total cfDNA concentration in cancer patients, we also observe an increase in the correlation between cfDNA concentrations originating from leukocytes and non-leukocyte sources, from weakly (anti-)correlated in healthy individuals, to significant positive correlation in cancer patients (Fig. 4a).

**Figure 4.**
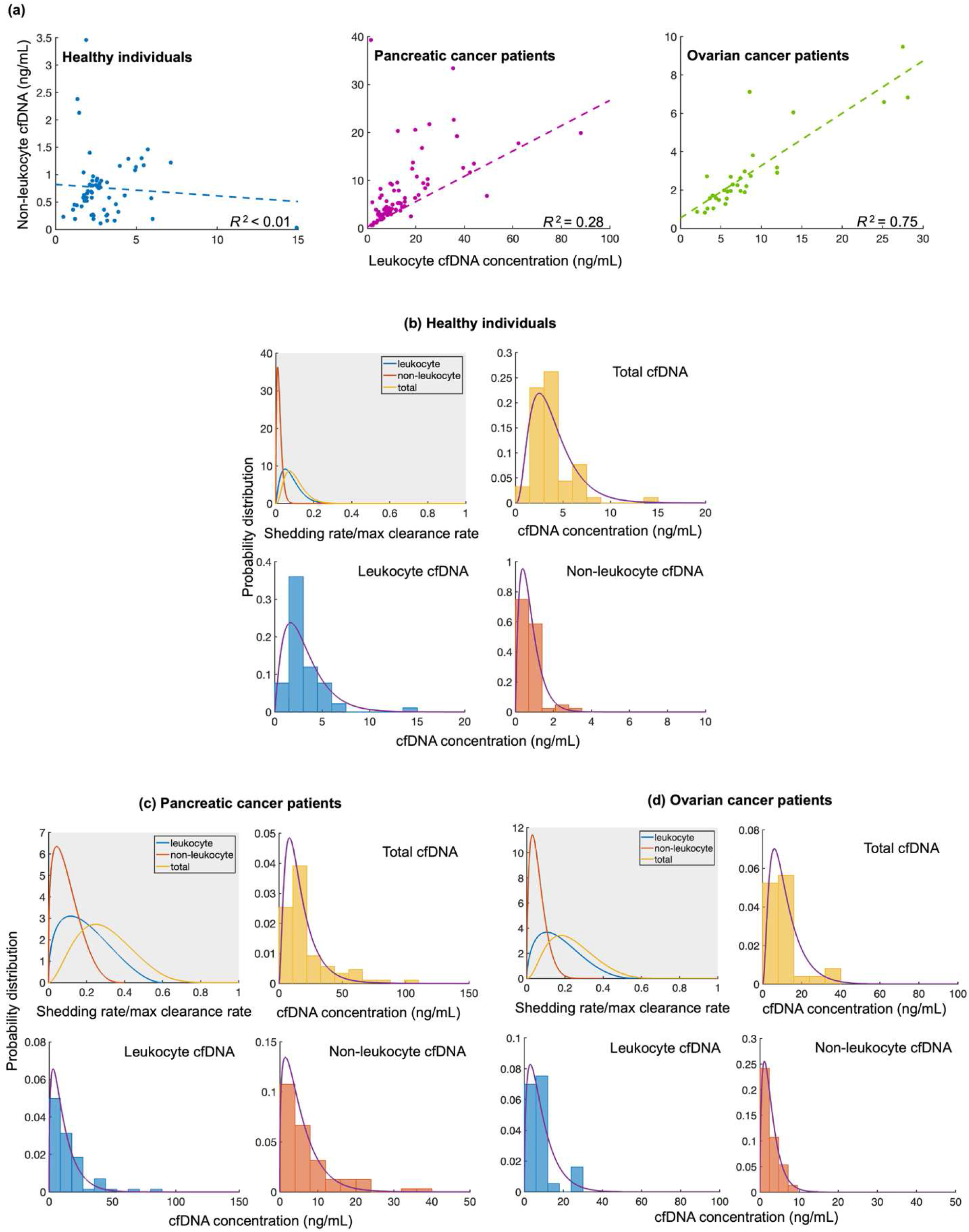
Correlation and distributions of the leukocyte, non-leukocyte and total cfDNA concentrations. **(a)** Correlation between leukocyte and non-leukocyte concentrations, excluding outliers with high cfDNA concentration. Dots represent the pair of leukocyte and non-leukocyte cfDNA concentration for normal individuals and pancreatic and ovarian cancer patients from Mattox *et al*. [19] under a quadratic programming deconvolution algorithm employing Sun *et al*. [23] reference matrix. **(b), (c), (d)** Gray panel: Estimated leukocyte 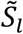, non-leukocyte 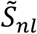 and total cfDNA 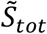 shedding rates normalized by the maximum clearance rate; 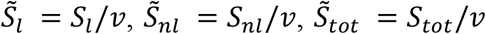, considered for the case of healthy individuals (b), pancreatic cancer patients (c), and ovarian cancer patients (d). White panels: Saturation model results for the distribution of leukocyte, non-leukocyte and total cfDNA concentrations are depicted by purple curves. Histograms represent data for cfDNA concentrations from Mattox *et al*. [19], under the same deconvolution method as for (a).

Stratifying by stage, we find correlations of 0.57–0.80 for stage I (*N*=13), 0.29–0.58 for stage II (*N*=60), and 0.33–0.51 for stage III (*N*=6) in pancreatic cancer, and 0–0.75 for stage I (*N*=8), 0.46–0.89 for stage II (*N*=5), and 0.33–0.88 for stage III (*N*=18) in ovarian cancer. These results indicate no consistent increase in correlation with stage; in fact, pancreatic cancer shows the highest correlation in stage I. Because stage-specific sample sizes are small, correlation estimates may exhibit high variability. To mitigate this, we performed a two-dimensional Kolmogorov–Smirnov (2D KS) test (Materials and Methods) on paired leukocyte and non-leukocyte cfDNA concentrations for each cancer type and deconvolution method. For samples of the same cancer type and deconvolution method but of different stages, the 2D KS test did not reject the null hypothesis that they originate from the same joint distribution, suggesting that cfDNA composition does not significantly change across stages I-III in pancreatic and ovarian cancer.

We formulate a mathematical model for cfDNA dynamics in order to obtain insight into the changes in cfDNA shedding or clearance that could lead to the observed multiplicative increase in total cfDNA concentration. The cfDNA dynamics model should also be able to explain the increase in correlation between cfDNA concentrations originating from leukocytes and non-leukocyte sources observed in cancer patients.

We first consider a model for cfDNA dynamics with linear clearance

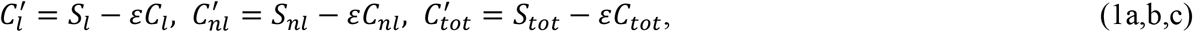

where *C*_*l*_, *C*_*nl*_ are the cfDNA concentrations and *S*_*l*_, *S*_*nl*_ are the rates of cfDNA shedding from leukocytes and non-leukocyte sources respectively. *C*_*tot*_ = *C*_*l*_ + *C*_*nl*_ and *S*_*tot*_ = *S*_*l*_ + *S*_*nl*_ are the total cfDNA concentration and the total cfDNA shedding rate. In Eqs. (1), *ε* denotes the rate of cfDNA clearance; it is related to cfDNA half-life *t*_*1/2*_ as ε=*ln* (*2*)*/t*_*1/2*_. We assume cfDNA dynamics reach a quasi-steady state between shedding and clearance. This assumption is supported by the rapid cfDNA clearance (half-life 15 minutes to 2.5 hours [5,6,13–15]), compared to tumor growth and therapy-driven changes in shedding rates, which occur over days to weeks, as reported in longitudinal studies of circulating tumor DNA (ctDNA) dynamics during cancer progression and therapy response [25–28]. Assuming quasi-steady state is reasonable for modeling purposes and has also been adopted in previous mechanistic studies [13,27]. Moreover, as noted earlier, two-dimensional Kolmogorov–Smirnov tests on the Mattox et al. [19] dataset indicate no significant differences in cfDNA levels or composition from leukocyte and non-leukocyte sources across stages I–III for pancreatic and ovarian cancer. Given the rapid clearance and the absence of stage-specific variation, it is reasonable to assume that cfDNA levels are at quasi-steady state. Under this assumption, the cfDNA concentrations calculated from the linear model (1) are

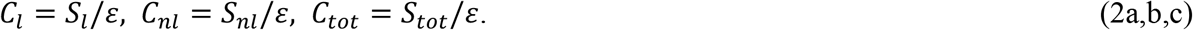

Under Eqs. (2) of the linear clearance model, the observed increases in cfDNA concentrations from leukocytes (*C*_*l*_) and non-leukocyte sources (*C*_*nl*_) may originate from corresponding increases in cfDNA shedding rates or from a decrease in the clearance rate *ε*. From the data by Mattox *et al*. [19], we calculate the magnitude of the multiplicative increases in the distributions of leukocyte and non-leukocyte cfDNA: 3.73 (3.15-4.45) and 7.99 (6.24-9.94) for pancreatic cancer, and 2.53 (2.03-3.37), 3.29 (2.39-4.49) for ovarian cancer. A decrease in cfDNA clearance rate should in principle increase both leukocyte and non-leukocyte cfDNA by the same factor. In pancreatic cancer, the multiplicative increase in cfDNA from non-leukocyte sources is significantly higher than from leukocytes. We deduce that the increased cfDNA concentration in the presence of pancreatic cancer does not originate from decreased cfDNA clearance only, but stems at least partly from increased cfDNA shedding from non-leukocyte and potentially also leukocyte sources.

We next investigated if the linear model of cfDNA dynamics can account for the observed increase in correlation between leukocyte and non-leukocyte cfDNA in cancer patients. We first note that a decrease in cfDNA clearance cannot explain increased correlation under the linear model (1). On the other hand, increases in cfDNA shedding from leukocyte and non-leukocyte sources may be correlated as they may both be affected by the presence of cancer. To capture this, we allow the multiplicative factors for leukocyte and non-leukocyte cfDNA shedding to vary across patients according to uniform distributions over the ranges of values determined previously from the KS test. We further assume that the multiplicative factors are perfectly linearly correlated, so that cancer presence increases shedding from both sources simultaneously. We provide an analytical calculation of the Pearson correlation coefficient for leukocyte and non-leukocyte cfDNA concentrations in cancer under the linear clearance model after applying multiplicative shedding factors to cfDNA concentrations from healthy individuals (Supplementary Text S2). From it we obtain the result that, even with variability in shedding increases across patients and perfect correlation between the shedding factors, the linear clearance model predicts the same correlation coefficient (from -0.29 to 0.20) between leukocyte and non-leukocyte cfDNA in cancer patients as in healthy individuals, for the four deconvolution methods. Therefore, the linear clearance model cannot easily account for the observed increase in cfDNA correlation in cancer patients.

As the linear model (1) is insufficient to explain the correlation observed in data, we consider a model with saturation in cfDNA clearance:

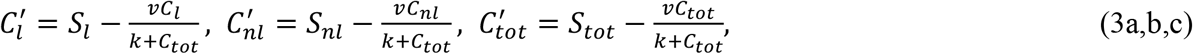

with *v* and *k* being positive constants. In Eq. (3c) for the total cfDNA concentration, clearance term is of Michaelis-Menten type [29]: for low cfDNA concentrations, the clearance term is approximately linear with respect to cfDNA concentration, with clearance rate *v/k*; as the total cfDNA concentration increases, the clearance term approaches *v*, which is the maximum cfDNA clearance rate. The Michaelis-Menten clearance term is able to model the finite capacity of Kupffer liver cells where most of cfDNA (70%-90%) is cleared from circulation [11,12]. In Eqs. (3a,b), the clearance rate for leukocyte and non-leukocyte cfDNA is the same and equal to the clearance rate *v/*(*k* + *C*_*tot*_) for the total cfDNA. This is based on the assumption that all cfDNA fragments are well mixed in blood circulation, and that all undergo the same clearance process, regardless of their tissue of origin. From Eqs. (3), the equilibrium values of leukocyte, non-leukocyte and total cfDNA concentrations in plasma are given by:

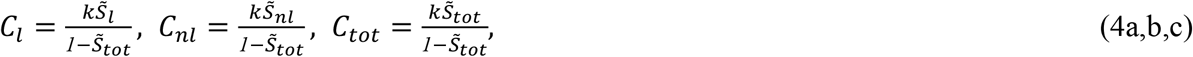

where 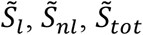 are the leukocyte, non-leukocyte and total shedding rates normalized by the maximum clearance rate; 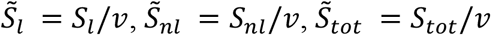. In our analysis, we impose that the distribution of the total normalized shedding rate 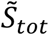 remains below 1. This reflects the physiological requirement that total cfDNA shedding should not exceed the maximum clearance capacity, as doing so would lead to unbounded cfDNA accumulation in circulation. Furthermore, normalization provides a dimensionless scale (0–1), which enables direct comparison between shedding distributions in healthy and cancer conditions and facilitates clear interpretation and quantitative assessment of how cfDNA shedding from healthy tissue shifts in the presence of stage I-III cancer (Fig. 4). Under the saturation in clearance model (3), the correlation between leukocyte and non-leukocyte cfDNA concentrations is positive and increases as the average normalized shedding rate from leukocytes and/or non-leukocyte sources is increased (Eq. S13 in S3 Text).

To illustrate the ability of the saturation model (3) to explain the observed cfDNA dynamics, we determine model parameters that allow good agreement with observed data. First, by considering a weak correlation between leukocyte and non-leukocyte cfDNA concentrations in healthy individuals, we can estimate the model parameter *k*, as well as the mean values and correlations of variation (CV -standard deviation over average) of normalized cfDNA shedding rates from leukocytes and non-leukocyte sources (Materials and Methods). We choose rescaled Beta distributions to model the normalized shedding rates, in order to ensure that the maximum total shedding rate is less than maximum clearance rate. We estimate the values of distribution parameters using the estimated mean values and coefficients of variation for the normalized shedding rates from leukocytes and non-leukocyte sources in healthy individuals. Substituting the estimate for *k* and the fitted distributions for 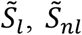 into Eqs. (4), we derive distributions for leukocyte, non-leukocyte and total cfDNA concentrations (Materials and Methods) that are in agreement with the data from Mattox *et al*. [19] for healthy individuals (K-S test *p*-value>0.05 for all distributions, Fig.4b). Furthermore, the correlation between leukocyte and non-leukocyte cfDNA concentrations predicted by the model is weak, ∼0.1, in agreement with the correlation -0.29 to 0.20 calculated for healthy individuals from Mattox *et al*. [19].

By considering a 2.5-fold increase in the average leukocyte shedding rate and a 5-fold increase in the average non-leukocyte shedding rate, saturation model (3) predicts distributions for leukocyte, non-leukocyte and total cfDNA concentrations that are in agreement with the data for pancreatic cancer patients (K-S test *p*-value>0.05 for all distributions, Fig.4c). Furthermore, it predicts a correlation coefficient 0.46 between leukocyte and non-leukocyte cfDNA concentrations, which is close to the median of the value range 0.31-0.58 calculated for pancreatic cancer patients in Mattox *et al*. [19] for different deconvolution methods. The aforementioned increases in the leukocyte and non-leukocyte shedding rates add up to a 3-fold increase in the average total shedding rate, and they account for a 4.4, a 3.6 and a 7.1-fold increase in the average total, leukocyte, and non-leukocyte cfDNA concentrations respectively. The resulting increases in cfDNA concentrations are greater than the respective increases in shedding rates, due to the saturation effect.

Similarly, by considering 2.2-fold and 3-fold increases in the average leukocyte and non-leukocyte shedding rates respectively, we obtain from model (3) distributions for all cfDNA concentrations that are consistent with the data for ovarian cancer patients (Fig. 4d). The correlation coefficient calculated from the model is 0.32, which is at the lower end of the value range 0.32-0.87 calculated for ovarian cancer patients in Mattox *et al*. [19] for different deconvolution methods.

### ROC curves for cancer detection using cfDNA level

Finally, in order to determine the implications of multiplicative cfDNA increase in cancer detection, we constructed the Receiver Operating Characteristic (ROC) curves for each cancer type from [20] individually (S8 Fig., S4 Table). ROC curves were generated using cfDNA concentration as the sole predictor, with thresholds scanned across its range, and constructed using 10-fold cross-validation with bootstrap resampling (Materials and Methods). Performance estimates were stable, as indicated by narrow confidence intervals obtained. We observe that, in cancer types with high multiplicative factors for cfDNA concentrations, such as ovarian, esophageal, stomach, and liver cancer, the area under the curve (AUC) approaches 1, being equal to or greater than 0.9. For lung, colorectal and stomach cancers where sufficient numbers of stage I cases were available, ROC curves and performance metrics were generally similar to that of the full cohort, indicating that cfDNA concentration retains its predictive value in early-stage disease. For ovarian cancer, AUC improved when considering only stage I cases, but this result should be interpreted with caution due to the small sample size (9 patients). Across all cancer types and stages, negative predictive value (NPV) remained consistently high (>0.85), underscoring the reliability of cfDNA concentration for ruling out disease even in early-stage cancers.

## Discussion

Our results indicate that cfDNA shed from healthy tissue exhibits a multiplicative shift towards higher concentrations in the presence of early-stage cancer; this increase is quantifiable and ranges from 1.3-fold in lung cancer to 12-fold increase in liver cancer. The agreement between the cancer-type specific multiplicative factors for total cfDNA concentration, and the concentration of circulating mutant DNA not originating from tumor, demonstrates the consistency of our finding.

We categorized cfDNA mutations in cancer patients as tumor or non-tumor derived based on whether the mutation detected in plasma was also detected in the tumor. Mutations detected in plasma but not in the tumor could still originate from the tumor if the biopsy missed regions harboring these mutations, and large-scale cfDNA clearance could further reduce plasma MAF relative to tumor MAF. While these possibilities cannot be excluded for individual cases, their overall impact appears limited:

When comparing plasma MAF distributions across three groups -mutations detected in plasma of healthy individuals, mutations detected in plasma but not in the tumor of cancer patients, and mutations detected in both plasma and tumor-we find that mutations not detected in the tumor cluster closely with mutations from healthy individuals (Kolmogorov–Smirnov test, p > 0.05; S9 Fig.). This concordance suggests that most plasma mutations not detected in the tumor indeed originate from non-tumor, healthy tissue.

Published data reinforce this interpretation: tumor-derived ctDNA in stage I–III cancers typically appears at 0.3–0.9% MAF [6,30], matching the median 0.3% MAF of the group of tumor-detected mutations from Cohen et al. [20]. On the other hand, cfDNA related to clonal hematopoiesis of indeterminate potential (CHIP), a major source of plasma mutations from healthy tissue, has MAF <0.1% [31,32]. In the Cohen et al. dataset [20], 92.5% of plasma mutation MAFs in healthy individuals and 87.4% of MAFs of plasma mutations not detected in the tumor of cancer patients were below 0.1%.

After we observed the multiplicative increase in cfDNA between healthy individuals and cancer patients, we asked about its potential diagnostic implications. Thus, we assessed the predictive performance of cfDNA concentration through receiver operating characteristic curves. For comparison purposes, we relate our findings to CancerSEEK, a multi-analyte blood test that integrates circulating-tumor DNA mutations and eight protein biomarkers [20]. CancerSEEK achieved an overall AUC of 91% (90–92%) when distinguishing cancer patients from healthy controls across the eight cancer types; at a fixed specificity >99%, the test reported a median sensitivity of 70% overall and 43% for stage I cancers, with variation by cancer type (e.g., 98% for ovarian, 100% for liver, and 33% for breast). In our analysis, cfDNA levels in plasma as the sole predictor produced AUC values above 0.90 for ovarian (0.901), stomach (0.918), esophageal (0.941), and liver cancers (0.951). For cancer types with sufficient number of stage I cases, we observed no major differences in sensitivity and specificity between all stages combined and stage I, suggesting that cfDNA concentration retains its discriminative ability even at early stages. Notably, our analysis shows that cfDNA concentration alone has limited predictive value in lung cancer. This is consistent with findings from the CancerSEEK study, which likewise reported lower sensitivity for lung cancer compared with most other cancer types. Differences in absolute specificity between CancerSEEK and our analysis reflect the fact that CancerSEEK optimized for ultra-high specificity, whereas our cutoff cfDNA levels balanced sensitivity and specificity. Overall, this comparison highlights that cfDNA concentration alone, without considering the mutations it harbors, can provide substantial diagnostic information, underscoring its potential as a simple and cost-effective component of liquid biopsy strategies for early cancer detection.

We also analyze an additional dataset [19] that reports the tissue of origin of cfDNA in healthy individuals and patients with pancreatic and ovarian cancer. From this dataset, we also observe another effect of cancer on cfDNA: a significant increase in the correlation between the cfDNA concentrations originating from leukocytes and non-leukocyte sources. Introducing simple models for cfDNA dynamics, we demonstrate that the linear clearance model, under which the increase in cfDNA levels originates from either a proportional increase in shedding rate or decrease in clearance rate, is insufficient to explain the observed increase in correlation. Rather, we show that the correlation between leukocyte and non-leukocyte cfDNA concentrations can be a manifestation of a saturation mechanism in cfDNA clearance. The saturation in clearance model is able to explain both the multiplicative increase in total cfDNA concentration and the increase in correlation, by considering increases in shedding rates that are smaller than the observed increases in cfDNA concentrations. The saturation effect is biologically plausible since cfDNA clearance occurs primarily via Kupffer cells in the liver that have a finite capacity [11,12]. For the cases of pancreatic and ovarian cancer where cfDNA composition was available [19], we observed a higher increase in cfDNA shedding from non-leukocyte sources compared to leukocytes. This disproportionate increase may reflect cancer-induced tissue stress and collateral damage in epithelial and stromal compartments, which are likely more vulnerable to inflammatory signaling and metabolic dysregulation than immune cells. These processes, including apoptosis, necrosis, and activation of stress-response pathways such as autophagy and DNA damage response, are documented in epithelial cells during cancer progression and can occur even in early-stage disease as part of adaptation to oncogenic and inflammatory stress [33–38]. We also note that, although NETosis can contribute to cfDNA release from leukocytes, its role is primarily associated with advanced cancer and metastatic processes rather than early-stage disease [39].

Highlighting the role of saturation effects in cfDNA clearance could contribute to existing mechanistic models, which often assume linear clearance [13,27]. This is relevant not only for early-stage cancer modeling, but also for monitoring and estimating tumor burden in later stages, where cfDNA kinetics inform treatment response and residual disease assessment.

We note that the reported cfDNA concentrations may differ significantly depending on the method used for the analysis of the samples. Employing a different methodology, Cohen *et al*. (2017) [40] report a median cfDNA concentration of 4.25ng/mL in healthy individuals, which is significantly different than the 2.83ng/mL reported in Cohen *et al*. (2018) [20]. However, we find that the multiplicative factor effect is also present between cfDNA concentrations from samples of healthy individuals from the two studies (multiplicative factor value 1.57 (1.34-1.79), S10 Fig.); this finding can be proven useful when cfDNA data from different studies are combined.

Finally, the cancer-specific multiplicative factor may be affected by the effects of therapy or anesthesia. Anesthesia has been reported to reduce the concentration of cfDNA ∼2.5-fold [41]. Cohen *et al*. (2017) [40] report cfDNA concentrations from 182 healthy individuals and 221 patients with stage I-III pancreatic cancer. As many pancreatic cancer patients in this study had their blood sample taken after administration of anesthesia, we expect that the multiplicative factor for pancreatic cancer is up to 2.5-fold lower compared to Cohen *et al*. (2018) [20], which excluded patients where anesthesia was known to be administered prior to sample collection. Indeed, we find that the multiplicative factor for pancreatic cancer from Cohen *et al*. (2017) [40] is 1.45 (1.30-1.59) (S11 Fig.). This factor is ∼2 times lower than the multiplicative factor for pancreatic cancer in Cohen *et al*. (2018) [20] data, as expected.

## Materials and Methods

### Patient data

We analyzed cfDNA concentrations (ng/mL) in plasma from healthy individuals and patients with stage I– III cancer reported by Cohen *et al*. [20]. The original dataset included 812 healthy individuals and 959 cancer patients across eight cancer types (lung, breast, colorectal, pancreatic, ovarian, esophageal, stomach, and liver). To ensure methodological consistency, we excluded 181 healthy individuals and 46 pancreatic cancer patients that were processed in a previous study using a different protocol [22], leaving 631 healthy individuals and 913 cancer patients for analysis (Table S1a). Among the 631 healthy individuals, 572 (90.6%) had circulating mutant DNA (cmDNA) detected in plasma and were included in comparative analyses, while 59 (9.4%) without detectable mutations were excluded from analyses involving cmDNA. Cohen *et al*. [20] also report the concentration (fragments/mL) and MAF% of the circulating mutant DNA (cmDNA) of the top DNA mutation detected in plasma for the same population of healthy individuals and cancer patients. The top mutation is chosen as the mutation with the highest omega score -a measure of statistical significance. Omega score is calculated in Cohen *et al*. [20] for every mutation in each sample based on the comparison between the mutation MAF and the reference MAF distributions among the healthy individuals and cancer patients in training sets. The panel for mutation detection from plasma includes 61 amplicons, each querying an average of 33 base pairs within one of the 16 cancer driver genes. Genes included in the panel and the number of amplicons for each gene are: *NRAS* (2), *CTNNB1* (2), *PIK3CA* (4), *FBXW7* (4), *APC* (2), *EGFR* (1), *BRAF* (1), *CDKN2* (2), *PTEN* (4), *FGFR2* (1), *HRAS* (1), *KRAS* (3), *AKT1* (1), *TP53* (31), *PPP2R1A* (1), *GNAS* (1). Apart from the top mutation detected in plasma, mutations found in tumor are also reported for 70 patients with lung cancer, 116 with breast, 336 with colorectal, 15 with pancreatic, 7 with ovarian, 36 with esophageal, 47 with stomach, and 24 with liver cancer. For these patients, we summarize in Table S1c the number of cases in which the top mutation in plasma was also detected in the tumor or not.

### Kolmogorov-Smirnov test

Kolmogorov-Smirnov (K-S) test is a nonparametric test for continuous one-dimensional probability distributions that we employ to check whether two datasets may come from the same distribution [42], Sec. 1.3.5.16. More specifically, in the case of two datasets, we use the Kolmogorov-Smirnov test to check whether the multiplication of all elements of one dataset by the same deterministic factor may result in the two datasets originating from the same distribution.

For the Mattox et al. [19] dataset, which includes paired leukocyte and non-leukocyte cfDNA concentrations are available, we also performed a two-dimensional Kolmogorov–Smirnov (2D KS) test on two-dimensional samples, each one consisting of leukocyte cfDNA concentration and non-leukocyte cfDNA concentration for every cancer patient. The test was used to compare the joint distributions of these paired concentrations across cancer stages. The 2D KS test is a nonparametric method that checks whether two 2D samples originate from the same bivariate distribution [43].

### Multiplicative factors for cfDNA/cmDNA

We determine the value of multiplicative factor *α* for which we obtain the minimum Kolmogorov distance between the empirical CDF for *α* times the sample of cfDNA concentration in healthy individuals with mutations detected in plasma, and the empirical CDF for the sample of cfDNA concentration in patients of each cancer type. We report the value of the multiplicative factor *α*, and the *p*-value of K-S test between the sample of cfDNA concentration in healthy individuals with mutations detected in plasma multiplied by *α*, and the cfDNA concentration in patients of each cancer type (Table S2).

We also report the value range of the multiplicative factor *α*, for which the K-S test does not reject, at 5% significance level, the hypothesis that cfDNA data for cancer patients and the cfDNA data for healthy individuals multiplied by the respective factor come from the same distribution. We also depict the P-P plots [42], Sec. 1.3.3.22. P-P plots are constructed by plotting the values of the empirical cumulative distribution functions (CDFs) of the two datasets for the same value of cfDNA concentration as a pair on the (x,y) plane. If all pairs stay close to the diagonal line from (0,0) to (1,1), this means that the two datasets follow the same distribution.

We calculate the multiplicative factors *α* between the cmDNA concentrations reported in healthy individuals, and the concentrations of cmDNA not shed from tumor in patients of the various cancer types. The top DNA mutation in plasma was not detected in the tumor for 43 patients with lung, 90 with breast, 205 with colorectal, 11 with pancreatic, 1 with ovarian, 18 with esophageal, 34 with stomach and 11 with liver cancer. The value of a resulting in the least Kolmogorov distance between the empirical CDFs, along with the 5% significance level ranges, are shown in Table S3; the respective P-P plots, which also stay close to the diagonal, are shown in Fig. S6. We exclude ovarian cancer from cmDNA concentration analysis, due to insufficient data. We also calculate the multiplicative factors *α* for cmDNA concentrations, excluding the samples with MAF less than i) 0.01%, ii) 0.015%, and iii) 0.02% (Table S3), since mutations detected in such low MAFs may be artifactual [22].

### Estimation of saturation model parameters and cfDNA shedding rates in healthy individuals

For the dataset of healthy individuals from Mattox *et al*. [19], we estimate parameter *k* of the cfDNA dynamics model (3) with saturation in clearance, as well as the averages 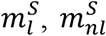 and coefficients of variation (standard deviation over average) 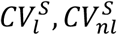 of normalized shedding rates 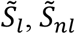 from leukocytes and non-leukocyte sources in the absence of cancer. First, we deduce from data a low correlation coefficient *ρ* = *0*.*1* between leukocyte and non-leukocyte concentrations in healthy individuals. Low correlation is predicted by the saturation model (3) for small average values of the normalized shedding rates, 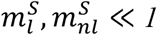, where Pearson’s correlation coefficient *ρ* between leukocyte and non-leukocyte cfDNA concentrations is approximated by the average total shedding rate (Eq. S19 in S3 Text):

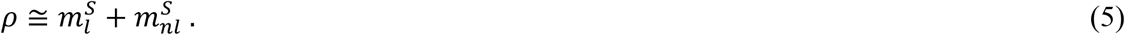

Furthermore, the average total cfDNA concentration, and the CV of leukocyte cfDNA concentration are expressed as (Eqs. S14c, S16a in S3 Text):

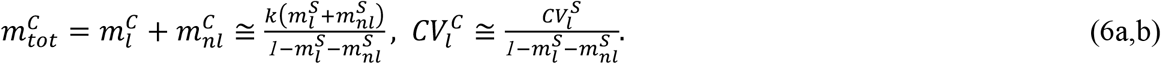

From Eqs. (5), (6a), and for a mean total cfDNA concentration 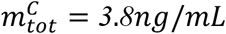 reported in healthy individuals by Mattox *et al*. [19], we estimate model parameter *k* to *34*.*2ng/mL*. Additionally, the ratios of the averages 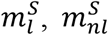, and the coefficients of variation 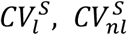 for shedding rates 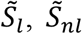 are approximated by the ratio of averages 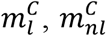 and 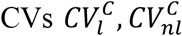 of the leukocyte and non-leukocyte cfDNA concentrations (Eqs. S15, S17 in S3 Text):

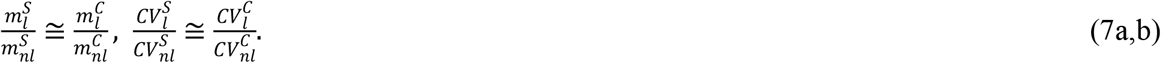

From Eqs. (5) and (7a), and for a reported 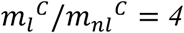 from the dataset of healthy individuals, we estimate the leukocyte and non-leukocyte average normalized shedding rates to 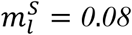 and 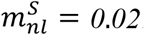 . Last, from Eqs. (6b), (7b) and for 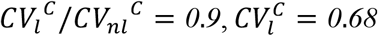 for healthy individuals as calculated from the Mattox *et al*. [19] dataset, we estimate 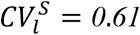 and 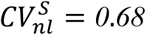 for the normalized shedding rates.

We use the estimated mean values and CVs to determine distributions for the normalized shedding rates. In order to ensure that the total normalized shedding rate is less than 1, we choose the rescaled Beta distribution for each of the normalized shedding rates

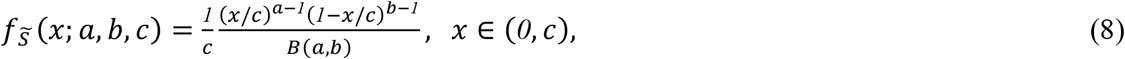

where *B*(*a, b*) is the Beta function. In Eq. (8), the parameter *c* represents the maximum normalized shedding rate for each source. We set *c* = 0.6 for leukocytes and *c* = 0.4 for non-leukocytes to reflect that, since cfDNA shedding occurs primarily during cell turnover [8,11], the ratio of leukocyte to non-leukocyte maximum shedding rates should approximate the ratio of their cell turnover rates, which is ∼1.5 [44]. The rescaled Beta distribution parameters *a, b* are determined via the mean *m*^*s*^ and coefficient of variation *CV*^*s*^ of each normalized shedding rate

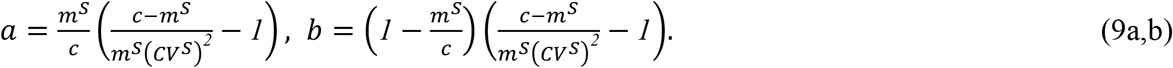

Assuming that the normalized shedding rates from leukocytes and non-leukocyte sources are independent, we determine via Eqs. (4a,b) the joint distribution of leukocyte and non-leukocyte cfDNA concentrations *C*_*l*_, *C*_*nl*_:

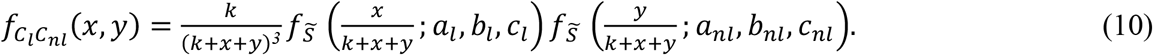

From the joint distribution (10), we determine the distributions for leukocyte, non-leukocyte and total cfDNA concentrations, as well as the correlation coefficient between leukocyte and non-leukocyte cfDNA concentrations.

### Construction of ROC curves

We computed Receiver Operating Characteristic (ROC) curves using cfDNA concentration as the predictor [45]. For each cancer type, the dataset was partitioned into *k* = 10 folds using cross-validation. In each iteration, 9 folds were used for training and 1 fold for testing. ROC curves were generated by plotting Sensitivity × 100% versus (1 − Specificity) × 100% across all possible thresholds, where:

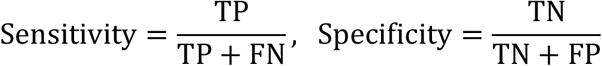

with TP, TN, FP, and FN denoting true positives, true negatives, false positives, and false negatives, respectively. Overall performance was summarized by the Area Under the Curve (AUC) and its 95% confidence interval, estimated using 1000 bootstrap resamples [46]. To quantify the variability in ROC curves arising from cross-validation fold partitioning, the 10-fold results were resampled with replacement for the 1000 bootstrap iterations. We created a grid of 200 fixed specificity values from 0 to 1, indexed by *j* = 1, …,200. For each bootstrap iteration, we computed the mean sensitivity 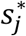 across folds at each specificity value from the grid. At every grid point *j*, we derived empirical distributions from the bootstrap sensitivity samples. The 95% confidence bands were then obtained as the 2.5th and 97.5th percentiles of these distributions:

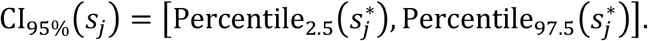

The optimal cutoff cfDNA level was selected by scanning all possible thresholds on the ROC curve and choosing the one that maximizes Youden’s *J* statistic, that balances sensitivity and specificity [45]:

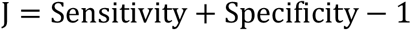

This threshold value was applied to the test fold to compute the positive predictive value (PPV), and negative predictive value (NPV):

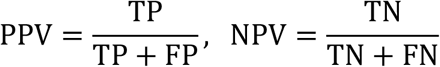

PPV represents the probability that a subject with a positive test truly has the disease, while NPV represents the probability that a subject with a negative test truly does not have the disease. PPV and NPV values reflect the prevalence of each cancer type in our study cohort. To quantify uncertainty in performance metrics we applied the same bootstrap resampling strategy used for AUC estimation. For each of the 1000 bootstrap iterations, the cross-validation folds were resampled with replacement, the optimal cutoff was re-estimated, and the metrics were recomputed. This generated empirical distributions for Sensitivity, Specificity, PPV, and NPV. The 95% confidence intervals for each metric were obtained as the 2.5th and 97.5th percentiles of the corresponding bootstrap distributions:

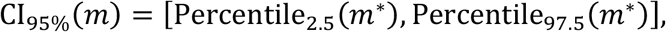

where *m* ∈ {Sensitivity, Specificity, PPV, NPV} and *m*^∗^ denotes bootstrap samples of metric *m*.

## Acknowledgements

The authors thank Bert Vogelstein for helpful comments on the manuscript.

## Funding

National Science Foundation grant DMS-2045166 (KM, IB)

Pacific Institute for the Mathematical Sciences (KM).

## Author contributions

KM and IB designed and performed research and wrote the manuscript.

## Artificial Intelligence declaration

No AI technologies have been used in the creation of this manuscript.

## Data and materials availability

The data analyzed in this study were obtained from Supplementary Materials of

Cohen *et al*. (2018) [20] https://doi.org/10.1126/science.aar3247

Cohen *et al*. (2017) [40] https://doi.org/10.1073/pnas.1704961114

Mattox *et al*. (2023) [19] https://doi.org/10.1158/2159-8290.CD-21-1252

MATLAB code used to generate the results and figures of this work is publicly available at: https://doi.org/10.5281/zenodo.19026910

## Electronic Supplementary Material

### S1 Text. Correlation coefficients between leukocyte and non-leukocyte cfDNA

For the dataset and the four different deconvolution methods from Mattox *et al*. [19], we calculate the Pearson and Spearman correlation coefficients between the cfDNA concentrations from leukocytes and non-leukocyte sources, for healthy individuals

#### Correlation coefficients for healthy individuals

|  | Sun *et al.* [23]<br>QP | Sun *et al.* [23]<br>NNLS | Moss *et al.* [24]<br>QP | Moss *et al.* [24]<br>NNLS |
| --- | --- | --- | --- | --- |
| Pearson | 0.84 | 0.94 | 0.51 | 0.31 |
| Spearman | 0.20 | 0.50 | 0.02 | -0.25 |

From the discrepancy between Pearson and Spearman coefficients, we deduce that Pearson coefficient is affected by the presence of outliers; by excluding the three outliers with significantly higher cfDNA concentrations compared to the rest of the dataset, we calculate

#### Correlation coefficients for healthy individuals (excluding the three outliers)

|  | Sun *et al.* [23]<br>QP | Sun *et al.* [23]<br>NNLS | Moss *et al.* [24]<br>QP | Moss *et al.* [24]<br>NNLS |
| --- | --- | --- | --- | --- |
| Pearson | -0.07 | 0.20 | -0.16 | -0.29 |
| Spearman | 0.08 | 0.42 | -0.13 | -0.42 |

After excluding the three outliers, Pearson and Spearman coefficients are in better agreement, showing a weak correlation between leukocyte and non-leukocyte cfDNA concentrations, especially for the deconvolution methods using Sun et al. reference matrix. Calculating the correlation coefficients for the pancreatic and ovarian cancer patients excluding the possible outliers we obtain:

#### Correlation coefficients for pancreatic cancer patients

|  | Sun *et al.* [23]<br>QP | Sun *et al.* [23]<br>NNLS | Moss *et al.* [24]<br>QP | Moss *et al.* [24]<br>NNLS |
| --- | --- | --- | --- | --- |
| Pearson | 0.53 | 0.58 | 0.31 | 0.39 |
| Spearman | 0.72 | 0.73 | 0.47 | 0.44 |

#### Correlation coefficients for ovarian cancer patients (excluding the one outlier)

|  | Sun *et al.* [23]<br>QP | Sun *et al.* [23]<br>NNLS | Moss *et al.* [24]<br>QP | Moss *et al.* [24]<br>NNLS |
| --- | --- | --- | --- | --- |
| Pearson | 0.87 | 0.87 | 0.32 | 0.60 |
| Spearman | 0.85 | 0.85 | 0.18 | 0.29 |

We observe that both in pancreatic and ovarian cancer, the leukocyte and non-leukocyte cfDNA concentrations are strongly correlated. In our analysis we will employ the Pearson correlation coefficient since its calculation from both datasets and analytic distributions is straightforward.

### S2 Text. Correlation coefficient in cfDNA composition for the linear clearance model

The cfDNA concentration from leukocyte (*C*_*l*_) and non-leukocyte sources (*C*_*nl*_) in healthy individuals are random quantities, with mean values, variances and covariance

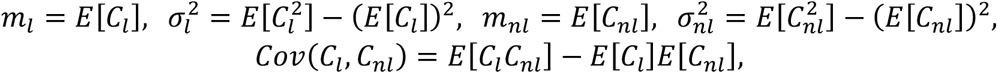

with E[·] being the expectation operator. Under the linear clearance model, *C*_*l*_ = *S*_*l*_*/ε, C_nl_* = *S*_*nl*_*/ε, C_tot_* = *S*_*tot*_ */ε*, multiplicative increases by factors *α*_*l*_, *α*_*nl*_ in leukocyte and non-leukocyte cfDNA shedding rates due to cancer presence result in a multiplicative increase of leukocyte and non-leukocyte cfDNA levels by the same factors

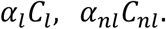

The multiplicative factors *α*_*l*_, *α*_*nl*_ in cancer are uncorrelated with the cfDNA levels *C*_*l*_, *C*_*nl*_ in healthy individuals. We assume that the multiplicative factors *α*_*l*_, *α*_*nl*_ vary across patients and follow uniform distributions over the ranges (*c*_*l*_, *d*_*l*_) and (*c*_*nl*_, *d*_*nl*_) respectively. These ranges are selected so that, when the healthy leukocyte and non-leukocyte cfDNA concentrations are multiplied by factors from these ranges, the resulting distributions satisfy the Kolmogorov–Smirnov (KS) test at the 5% significance level when compared to the corresponding cfDNA concentrations in cancer patients. Last, we assume that the multiplicative factors *α*_*l*_, *α*_*nl*_ are perfectly linearly correlated, meaning that cancer presence increases cfDNA shedding from both leukocyte and non-leukocyte sources simultaneously:

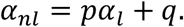

The constants p, q are determined as

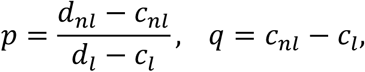

so that *α*_*nl*_ ∈ (*c*_*nl*_, *d*_*nl*_) for *α*_*l*_ ∈ (*c*_*l*_, *d*_*l*_).

To calculate the Pearson correlation coefficient between leukocyte *α*_*l*_*C*_*l*_ and non-leukocyte *α*_*nl*_*C*_*nl*_ cfDNA concentrations in cancer, we calculate their covariance

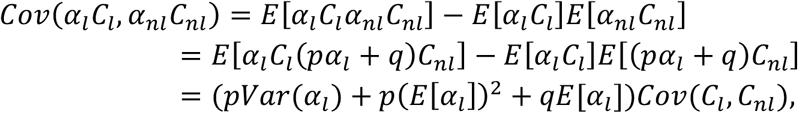

where *Var*(*α*_*l*_) is the variance of the multiplicative factor of cfDNA levels from leukocytes. We also calculate the variances

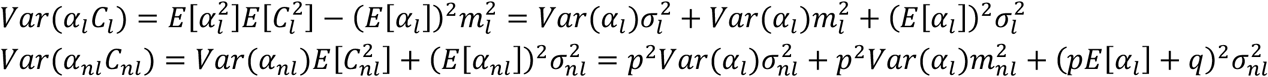

The Pearson correlation coefficient for *α*_*l*_*C*_*l*_, *α*_*nl*_*C*_*nl*_ reads

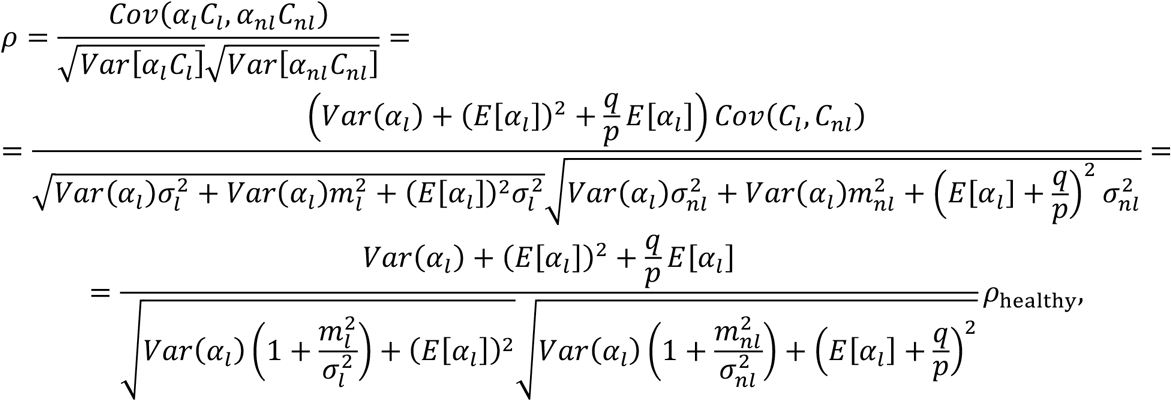

where 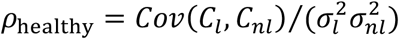 is the Pearson correlation coefficient between leukocyte and non-leukocyte cfDNA in healthy individuals. We have the ranges (*c*_*l*_, *d*_*l*_) = (3.15,4.45), (*c*_*nl*_, *d*_*nl*_) = (6.24,9.94) for pancreatic cancer and (*c*_*l*_, *d*_*l*_) = (2.03,3.37), (*c*_*nl*_, *d*_*nl*_) = (2.39,4.49) for ovarian cancer. Since we consider *α*_*l*_, *α*_*nl*_ to be uniformly distributed, we calculate the means and variances that appear in the above relation for correlation coefficient

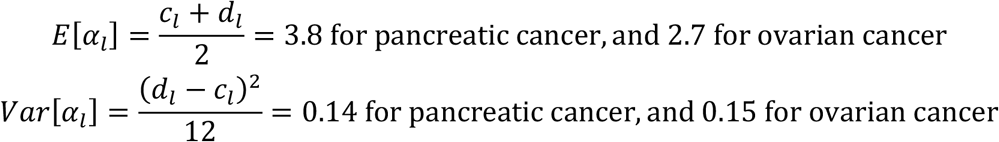

also

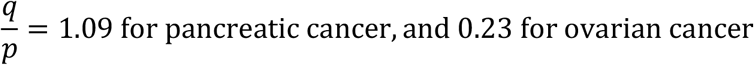

and from the data of healthy individuals, for the four deconvolution methods, we calculate

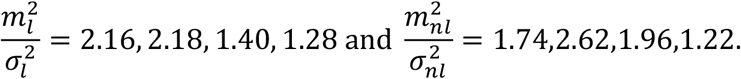

We observe that, for both cancer types, the mean dominates, *E*[*α*_*l*_] ≫ *Var*[*α*_*l*_], and also 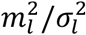 and 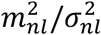 are of order one, so they do not offset the dominance. Thus, in the ratio multiplying p_healthy_, the terms dependent on *Var*[*α*_*l*_] can be omitted:

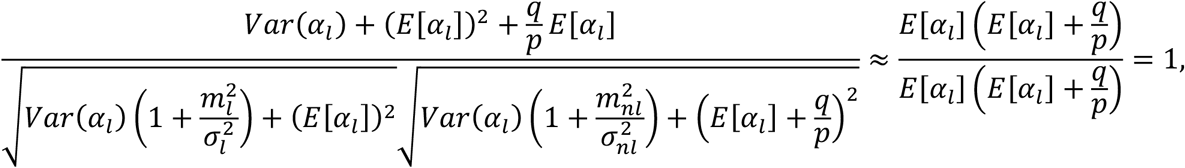

resulting in *ρ* ≈ *ρ*_heal*t*hy_ for both pancreatic and ovarian cancers. Indeed, by substituting the values of all parameters, for each cancer type separately and for each of the deconvolution methods, we get *ρ* from - 0.29 to 0.20, effectively coinciding with the Pearson correlation coefficient between leukocyte and non-leukocyte cfDNA concentrations in healthy individuals.

### S3 Text. Moment approximations for the model with saturation in clearance

For the saturation in clearance model,

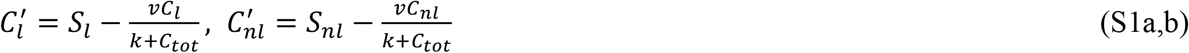

the equilibrium leukocyte, non-leukocyte and total cfDNA concentrations *C*_*l*_, *C*_*nl*_ and *C*_*tot*_ = *C*_*l*_ + *C*_*nl*_ are

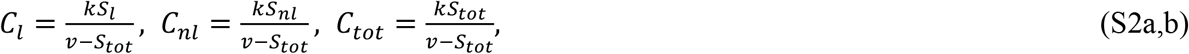

where *S*_*l*_, *S*_*nl*_ and *S* = *S*_*l*_ + *S*_*nl*_ are the leukocyte, non-leukocyte and total cfDNA shedding rates, *v* is the maximum cfDNA clearance rate, and *k* is the Michaelis-Menten parameter. By introducing the shedding rates normalized by the maximum clearance rate, 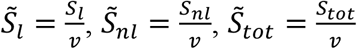 we express Eqs. (S2a,b) equivalently as

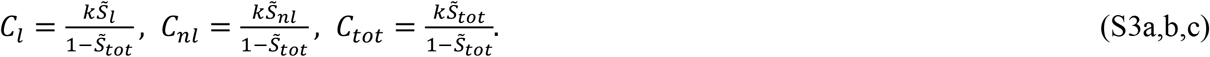

From Eqs. (S3a,b) we deduce that the saturation effect does not depend on the value of maximum clearance rate *v* per se; rather, it depends on total shedding rate as a percentage of *v*, i.e. 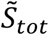 : For 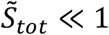, the saturation effect is negligible, with the concentrations approaching

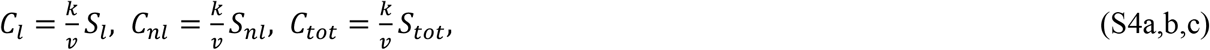

which are the equilibrium concentrations for the cfDNA model with linear clearance

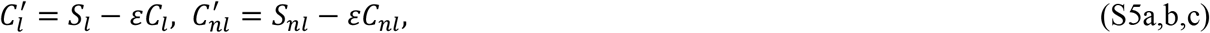

and clearance rate *ε* = *v/k*. Saturation in model (S1a,b) becomes more pronounced as the total shedding rate approaches the maximum clearance rate.

In our setting, we consider parameters *v, k* to be deterministic, while shedding rates *S*_*l*_, *S*_*nl*_ are independent random variables whose sum is less than *v*. We note that since parameter *k* enters formulas (S3a,b) for cfDNA concentrations as a deterministic multiplicative factor, average cfDNA concentrations depend on *k*,

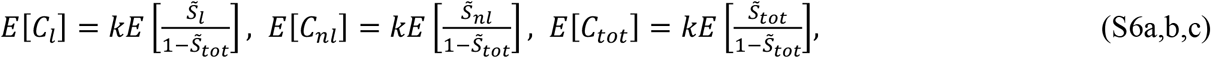

The Pearson correlation coefficient *ρ* and the coefficients of variation 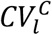 and 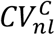 are *k* −independent:

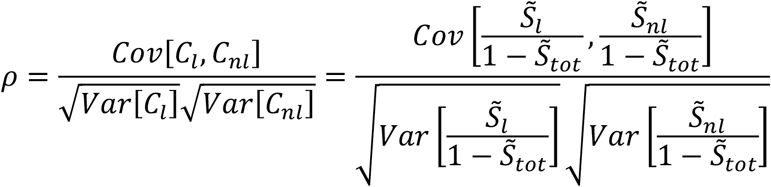

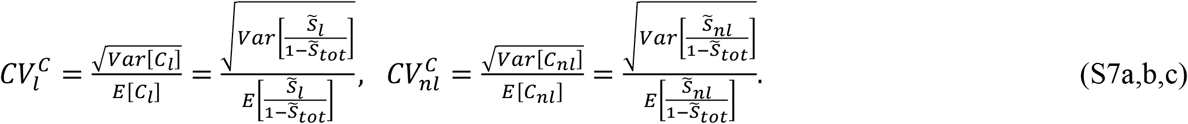

We approximate the moments of *C*_*l*_, *C*_*nl*_ by using Taylor expansions under the expectation operator, given that the moments of normalized shedding rates 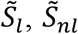 are known: The Taylor expansion for a function *f*(*x, y*) with two arguments is expressed as

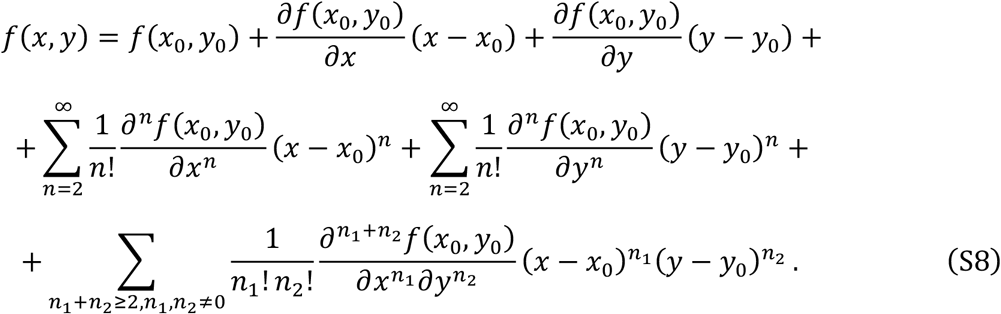

We choose 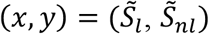, and 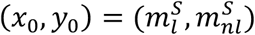 with 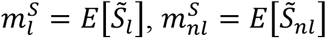 being the mean values, and take the expected value in both sides of Eq. (S8)

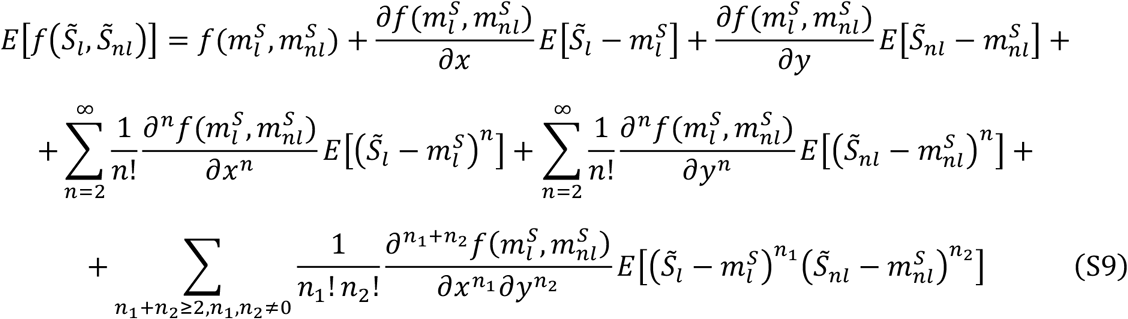

In the right-hand side of Eq. (S9), the terms containing the first-order partial derivatives of *f* are equal to zero, since 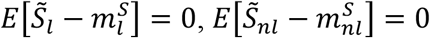. Furthermore, by assuming that normalized shedding rates 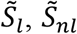 are independent, all cross moments in the last summation in the right-hand side of Eq. (S9) zero. Thus, we simplify Eq. (S9) to

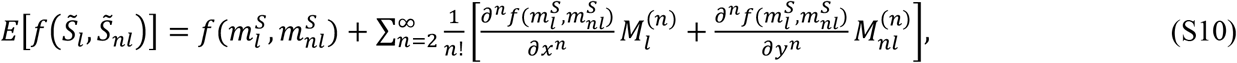

where 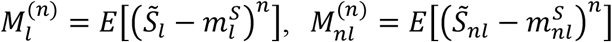 denote the centered moments of 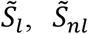 respectively. Using Eq. (S10), we calculate the series expansion for the first and second moments of *C*_*l*_, *C*_*nl*_, with 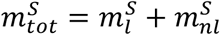 being the mean normalized total shedding rate:

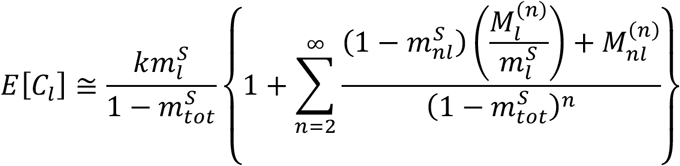

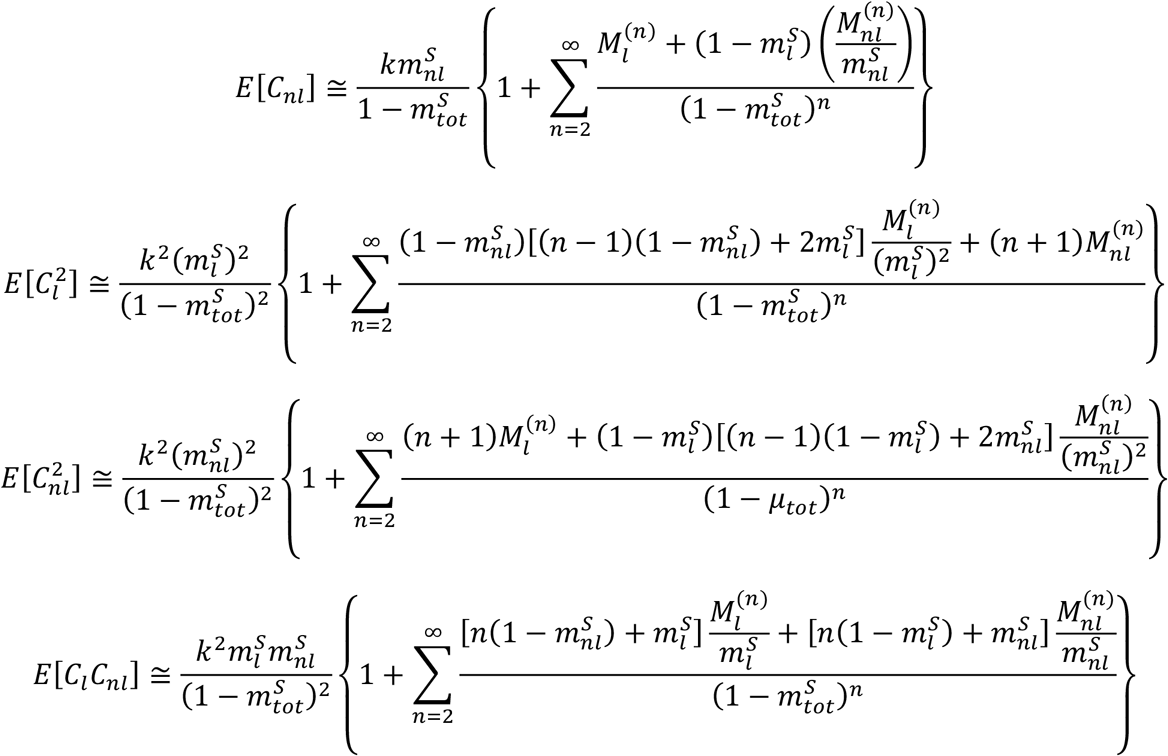

By keeping up to linear terms with respect to normalized shedding rates variances 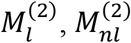 in the series expression above, we approximate the mean values, and centered second-order moments for cfDNA concentrations

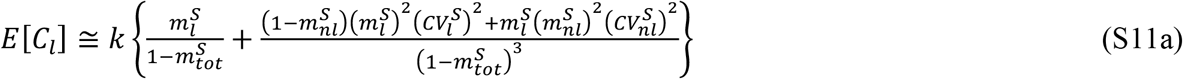

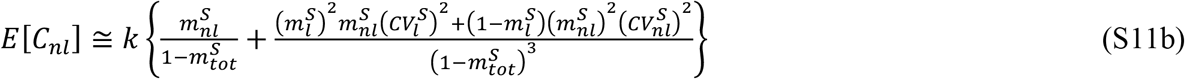

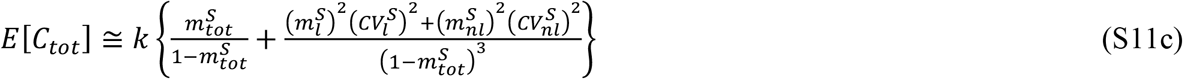

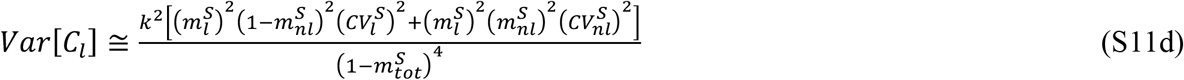

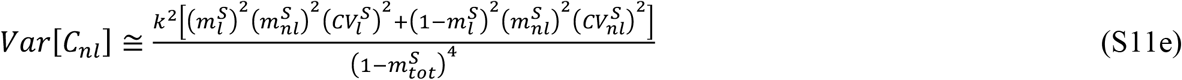

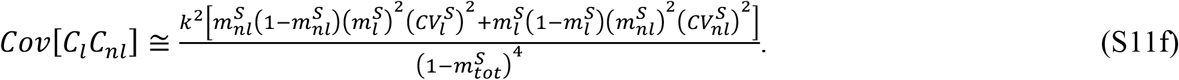

From Eqs. (S11) we approximate the coefficients of variation for the leukocyte and non-leukocyte cfDNA concentrations

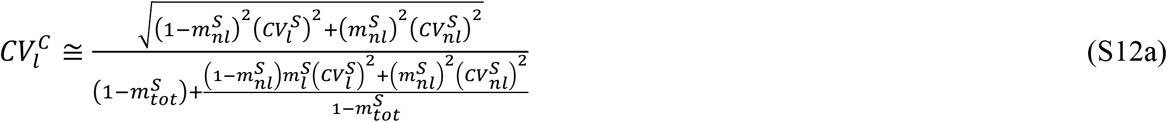

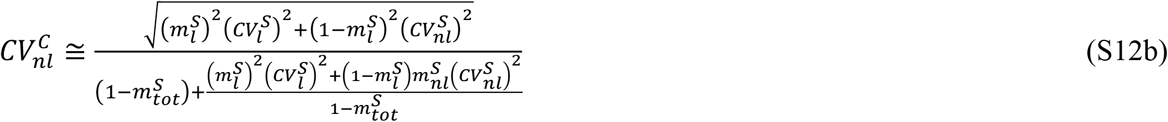

and the Pearson’s correlation coefficient between the leukocyte and non-leukocyte cfDNA concentrations

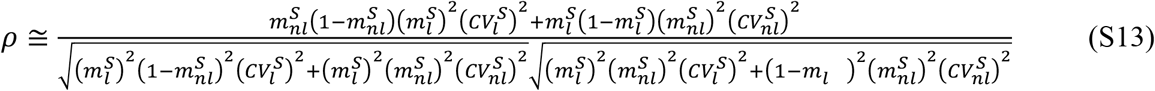

By calculating the derivative of *ρ* with respect to *m*_*l*_^*S*^ we get 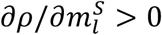. Since *m*^*S*^ and *m*^*S*^ appear in the right-hand side of Eq. (S13) in symmetric positions, 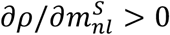 too. *Thus, the saturation in clearance model predicts increase in correlation between C*_*l*_ *and C*_*nl*_ *as the average normalized shedding rate from either leukocytes or non-leukocyte sources increases*.

#### Small shedding rates regime

For the case of small shedding rates 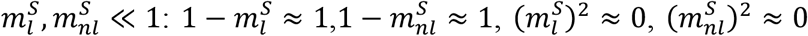.

Thus, Eqs. (S11) for mean cfDNA concentrations are further simplified to

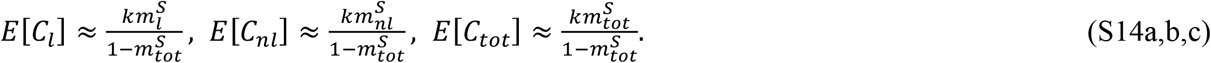

From Eqs. (S14) we also deduce that

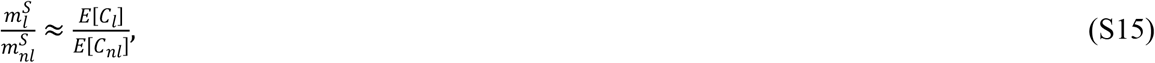

*i*.*e. the ratio of average leukocyte to non-leukocyte shedding rates is approximated by the ratio of average leukocyte to non-leukocyte cfDNA concentrations*.

Eqs. (S12) for the coefficients of variation are simplified to

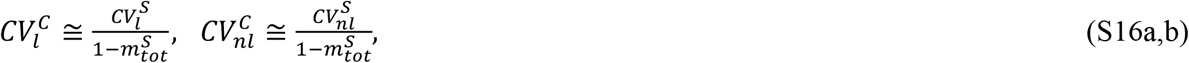

which implies that the saturation effect results in sightly increased coefficients of variation for cfDNA concentrations compared to the coefficients of variation of the respective shedding rates. From Eqs. (S16) we also deduce that

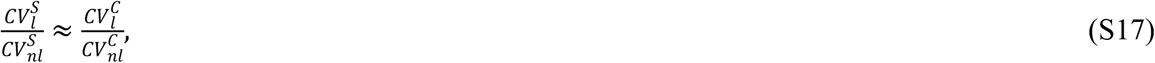

i.*e. the ratio of the coefficients of variation for leukocyte to non-leukocyte shedding rates is approximated by the ratio of the coefficients of variation for leukocyte to non-leukocyte cfDNA concentrations*.

Eq. (S13) for the correlation coefficient is simplified to

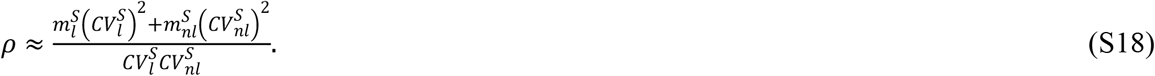

From Eq. (S18) we further deduce that, in the case where 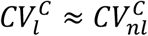, and thus 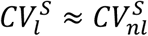 from Eq. (S17), *correlation coefficient is approximated by the average total normalized shedding rate:*

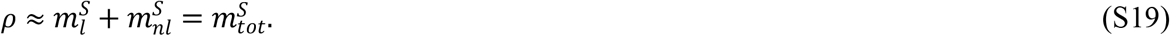

**S1 Figure.**
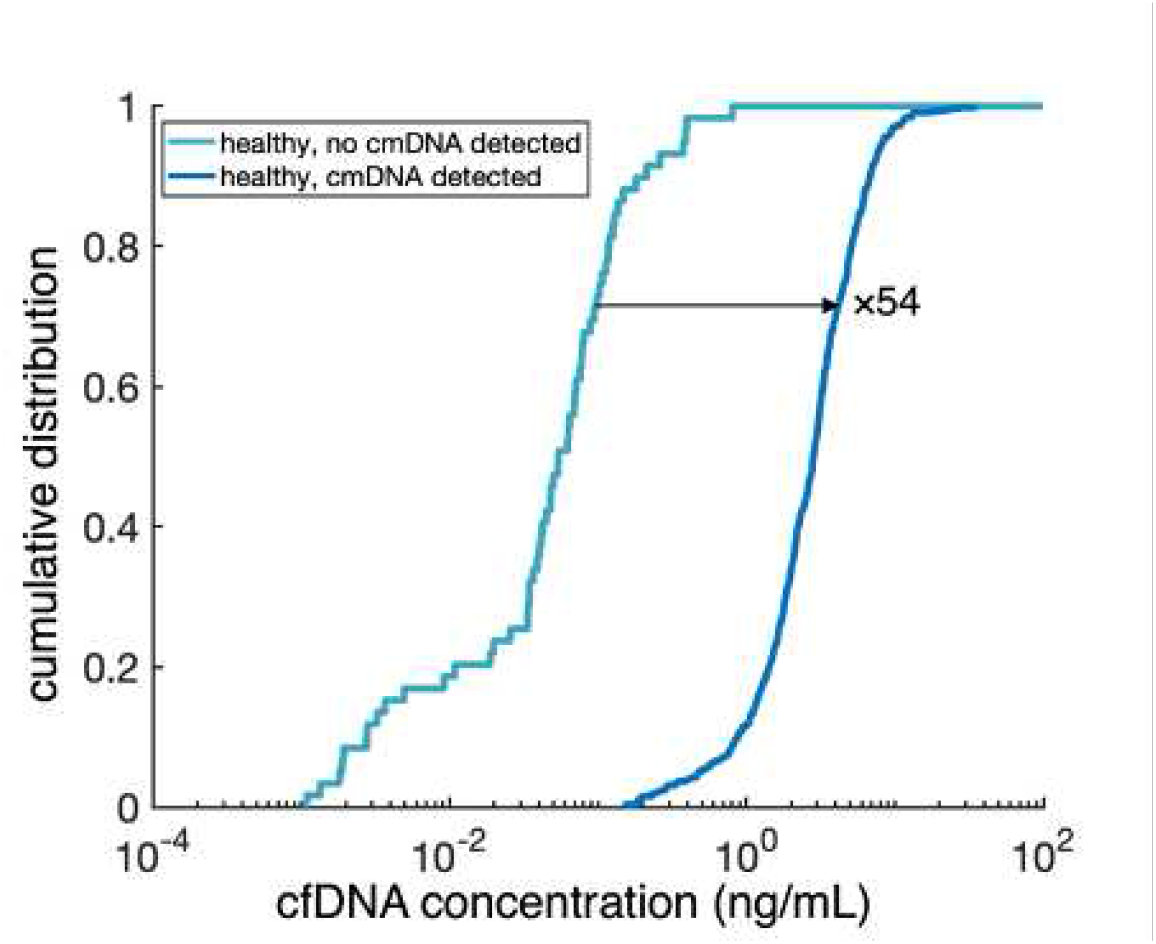
Multiplicative factor effect in cfDNA concentration between the 572 healthy individuals with cmDNA detected and the 59 healthy individuals without cmDNA detected.

**S2 Figure.**
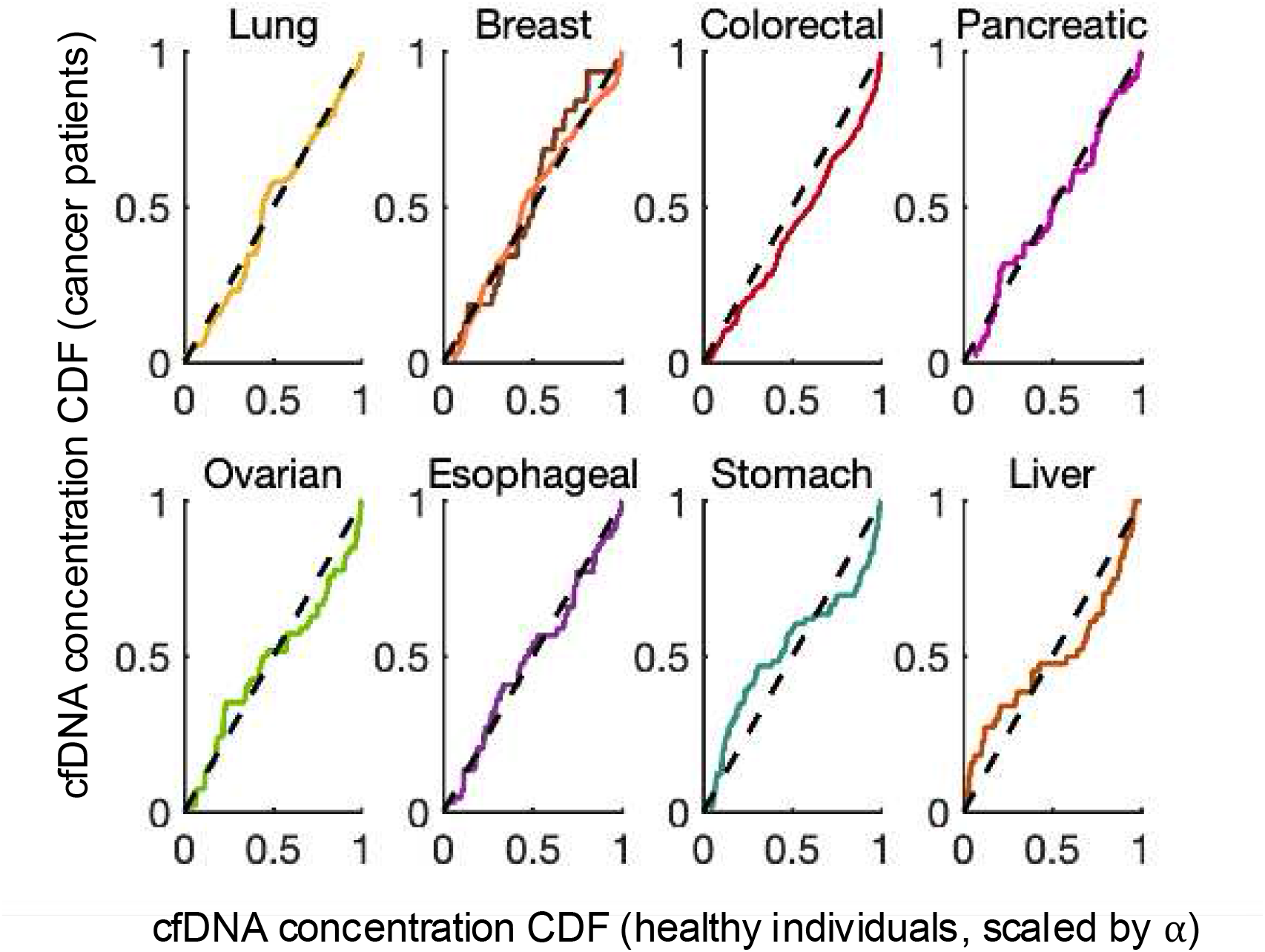
P-P plots for the effect of multiplicative factor *α* in cfDNA concentration between the 572 healthy individuals and the 103 patients with lung cancer, 209 with breast cancer, 385 with colorectal cancer, 47 with pancreatic cancer, 54 with ovarian cancer, 44 with esophageal cancer, 66 with stomach cancer, and 44 with liver cancer.

**S3 Figure.**
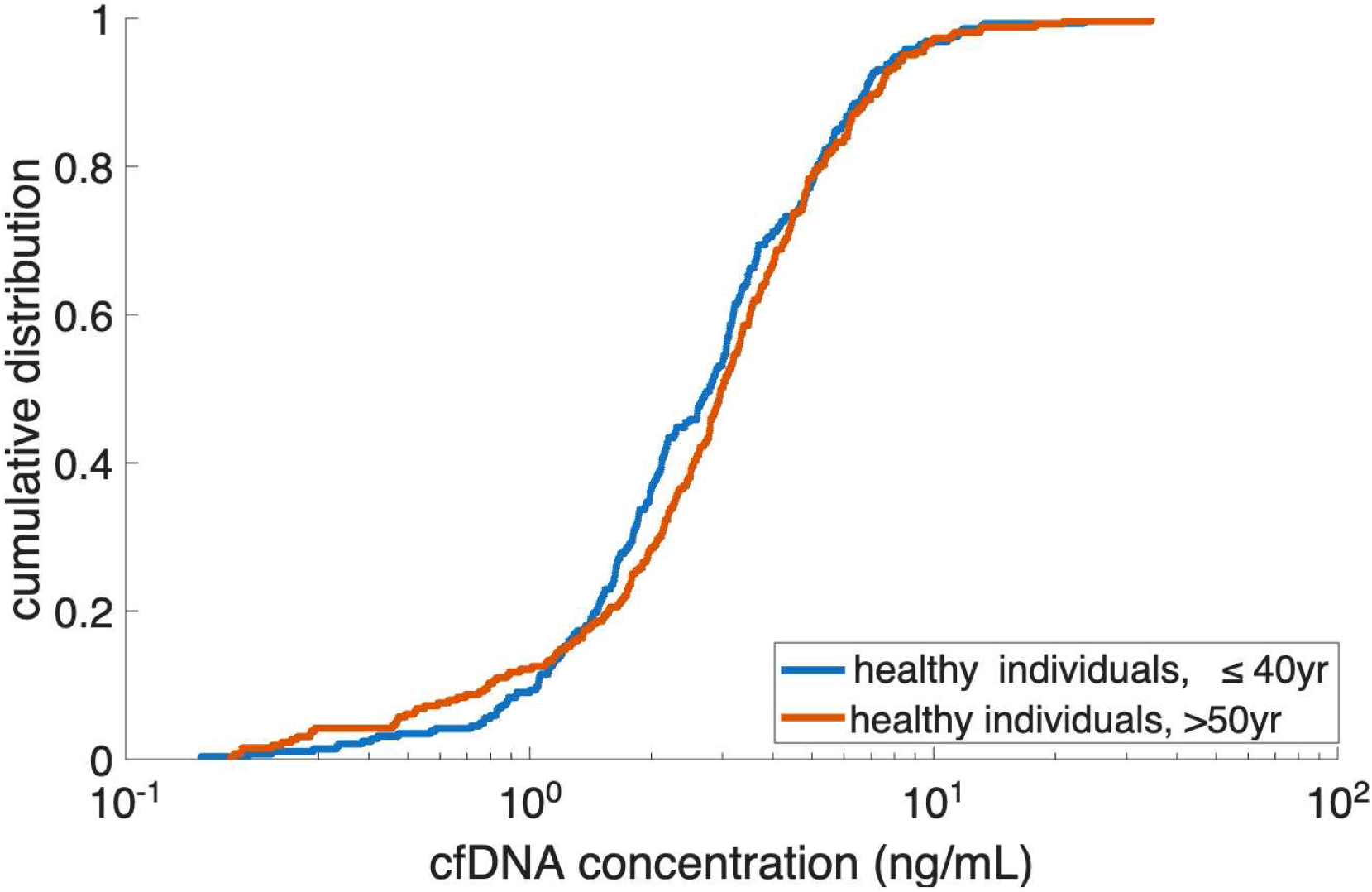
Cumulative distribution function for the cfDNA concentrations of the 288 healthy individuals under 40 years, and of the 263 healthy individuals over 50 years of age.

**S4 Figure.**
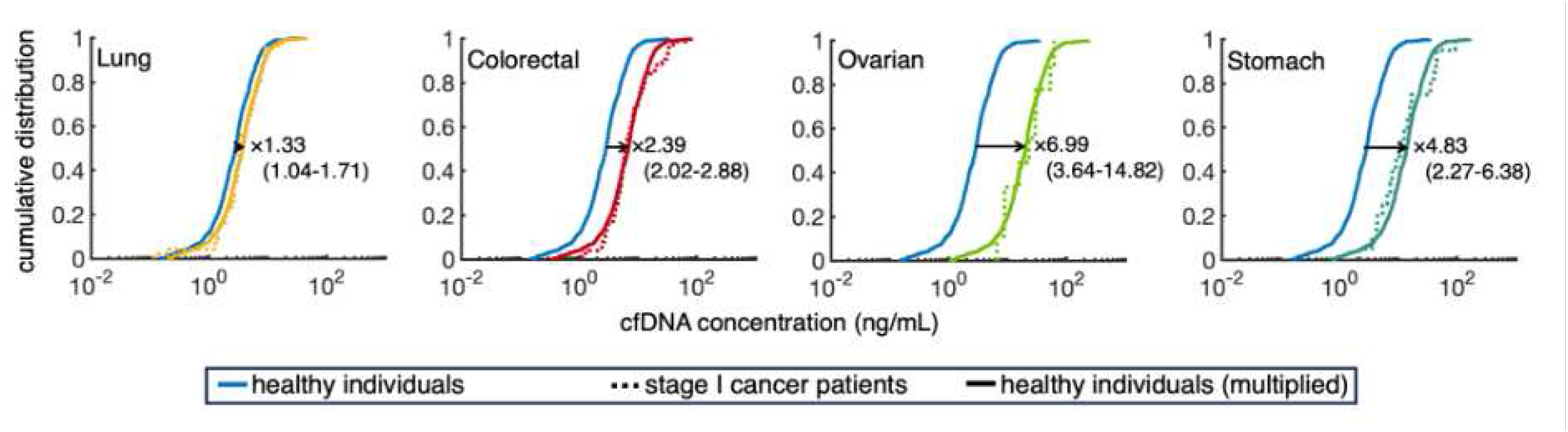
Quantifying the multiplicative increase in cfDNA concentration between the 572 healthy individuals and the 46, 76, 9 and 20 patients with stage I lung, colorectal, ovarian and stomach cancer respectively.

**S5 Figure.**
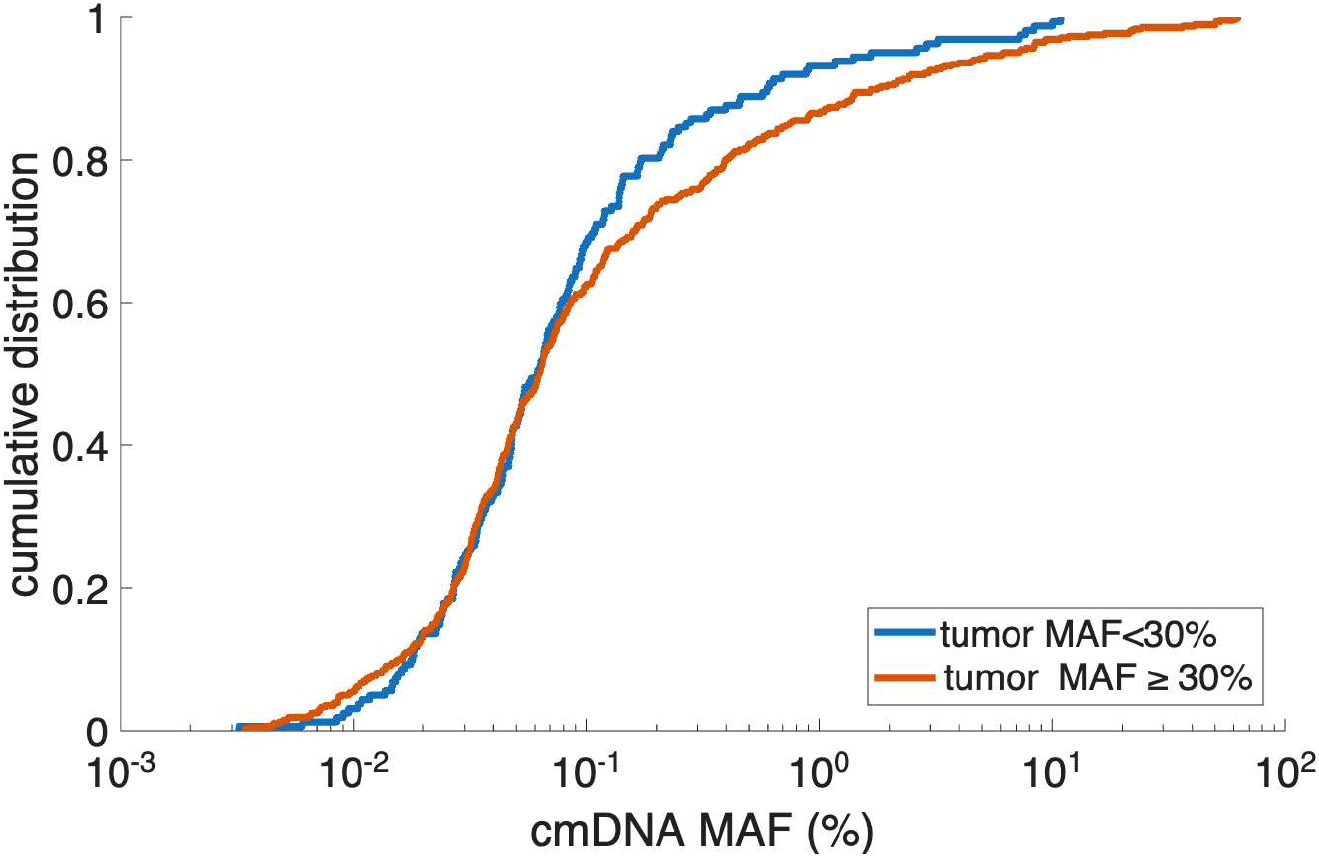
Cumulative distribution functions for the MAF of the top plasma mutation for the 162 cancer patients with all mutations with MAF<30% in the tumor (blue line), and for the 484 cancer patients with at least one mutation with MAF≥30% in the tumor (red line).

**S6 Figure.**
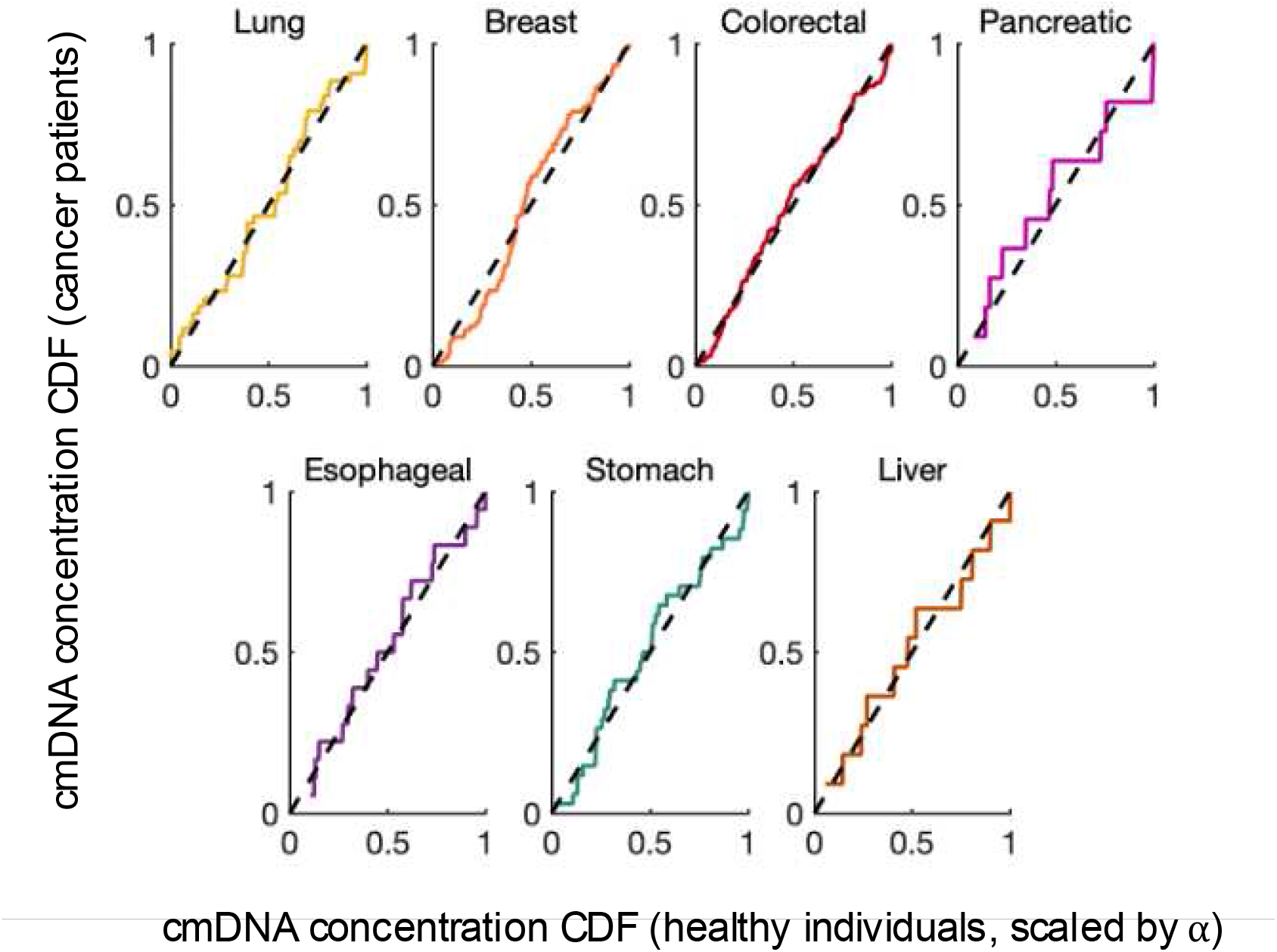
P-P plots for the effect of multiplicative factor *α* in cmDNA concentration between the 572 healthy individuals and the 43, 90, 205, 11, 18, 34, 11 patients with lung, breast, colorectal, pancreatic, esophageal, stomach and liver cancer respectively, with the top plasma mutation not detected in the tumor.

**S7 Figure.**
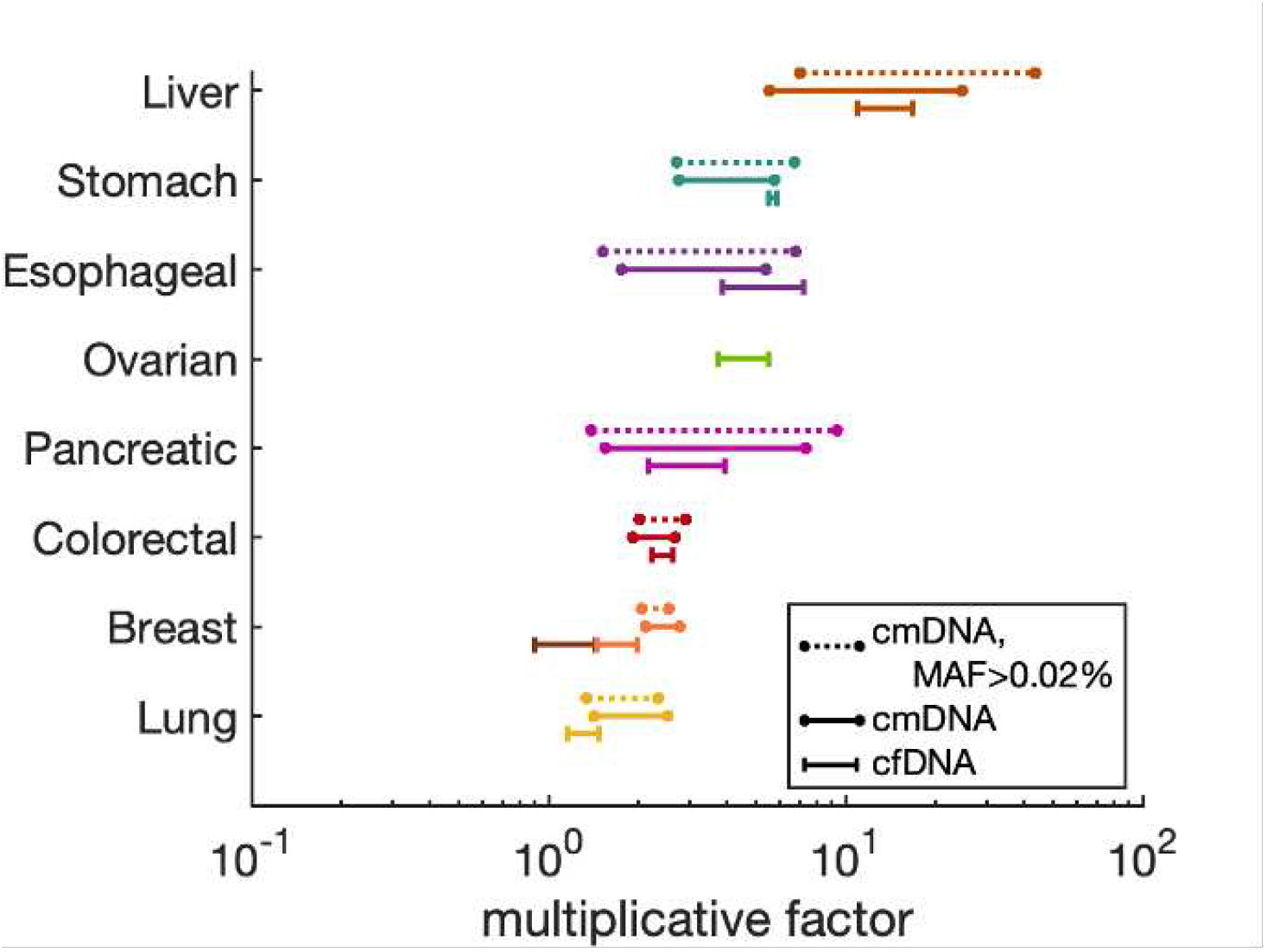
Multiplicative factor ranges for each cancer type, quantifying the increase in concentrations of total cfDNA in plasma, and of cmDNA not shed from cancer tissue, in the presence of cancer.

**S8 Figure.**
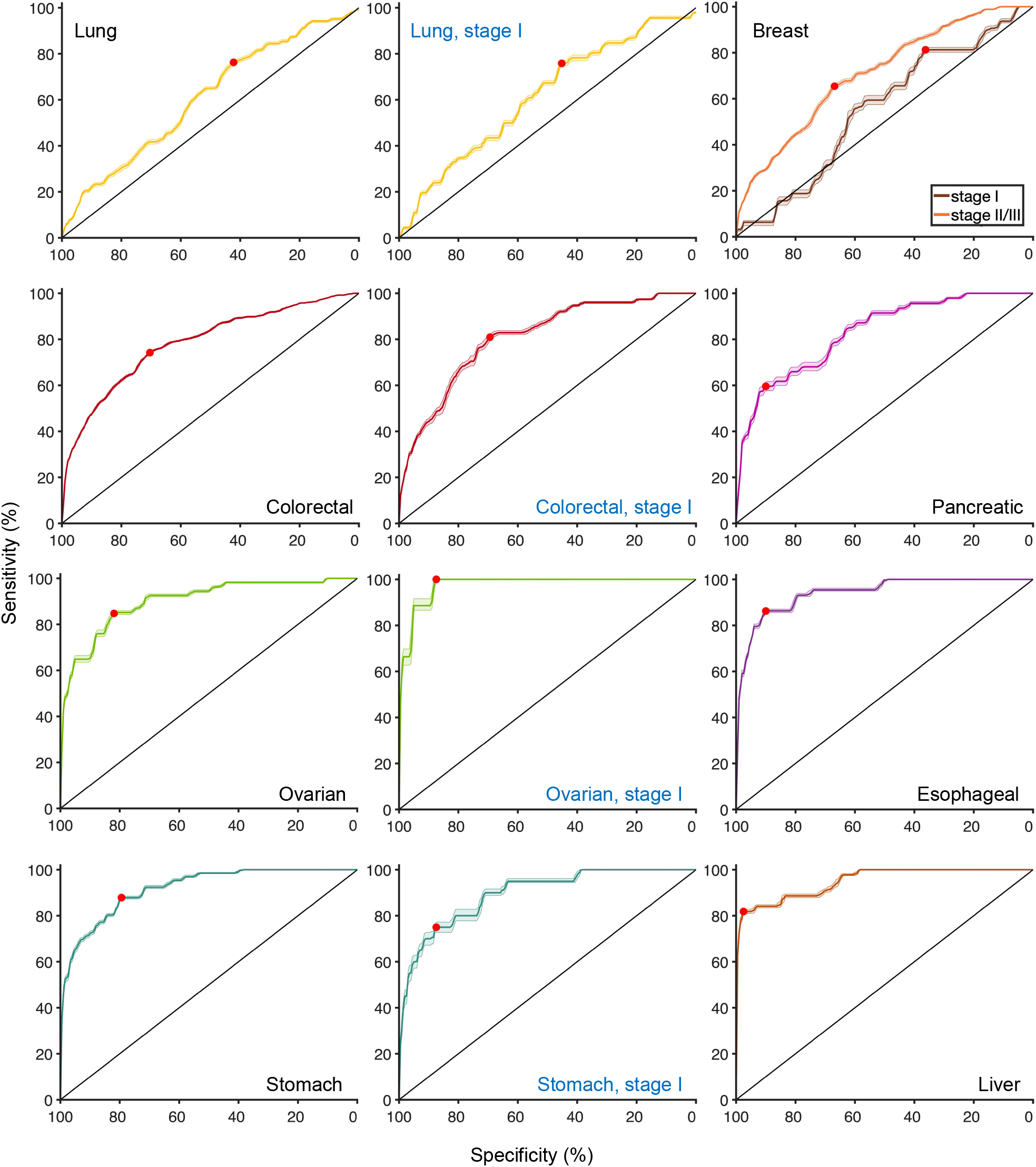
Receiver Operating Characteristic (ROC) curves with 95% confidence intervals for cfDNA-based discrimination between cancer patients and healthy controls. Solid lines indicate mean ROC curves across 10-fold cross-validation; shaded regions represent bootstrap-derived confidence intervals. Red dots mark the optimal cutoff determined by Youden’s J statistic on the mean ROC curve. Curves are shown for each cancer type (all stages) and for stage I subsets (lung, colorectal, ovarian, and stomach) to illustrate early detection performance. Pancreatic, esophageal, and liver cancers are not presented for stage I due to insufficient sample sizes.

**S9a Figure.**
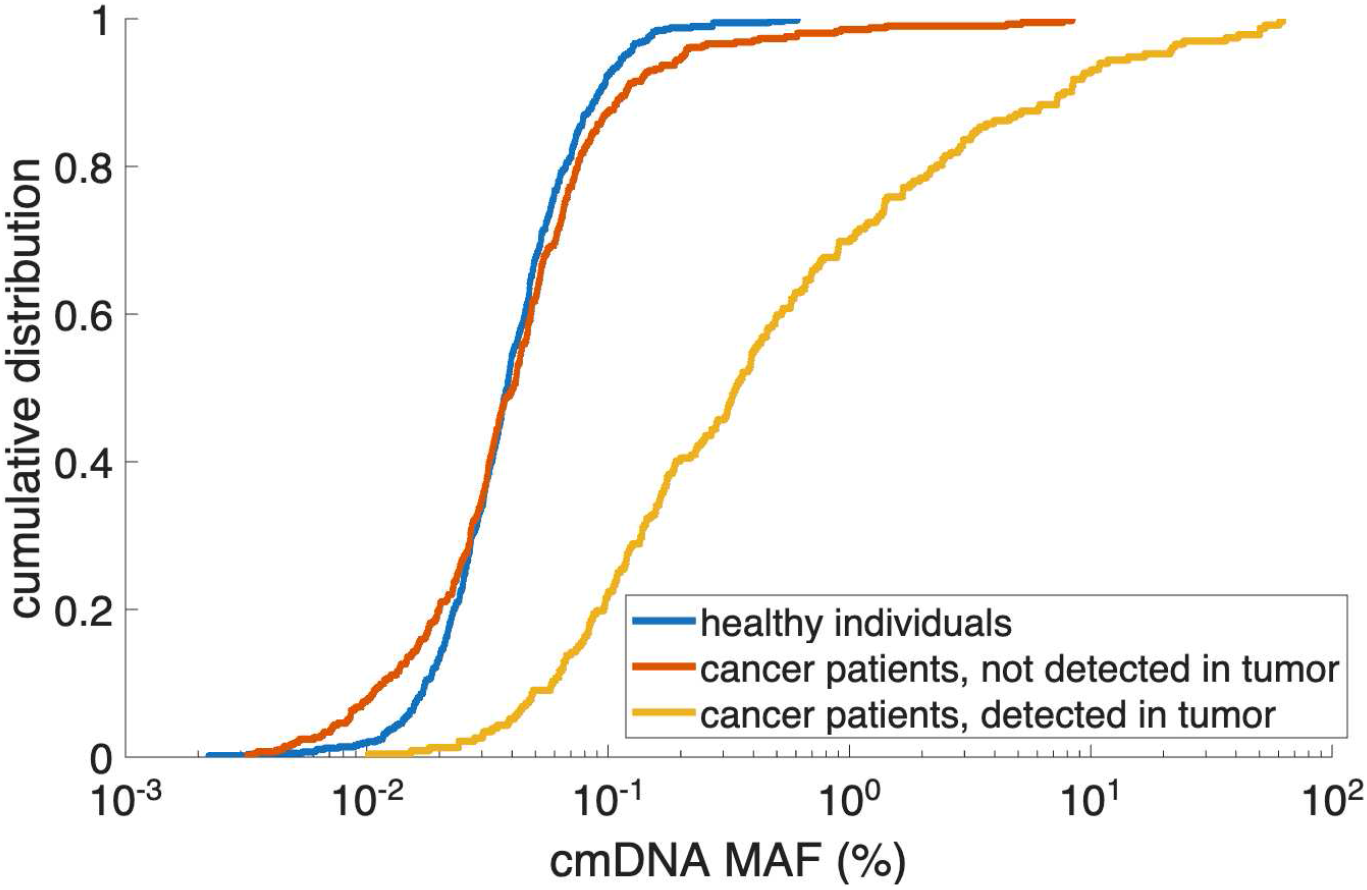
Cumulative distribution function for the MAFs of top plasma mutation in 572 healthy individuals, 413 cancer patients whose top plasma mutation was not detected in tumor, and for 232 cancer patients whose top plasma mutation was detected in the tumor.

**S9b Figure.**
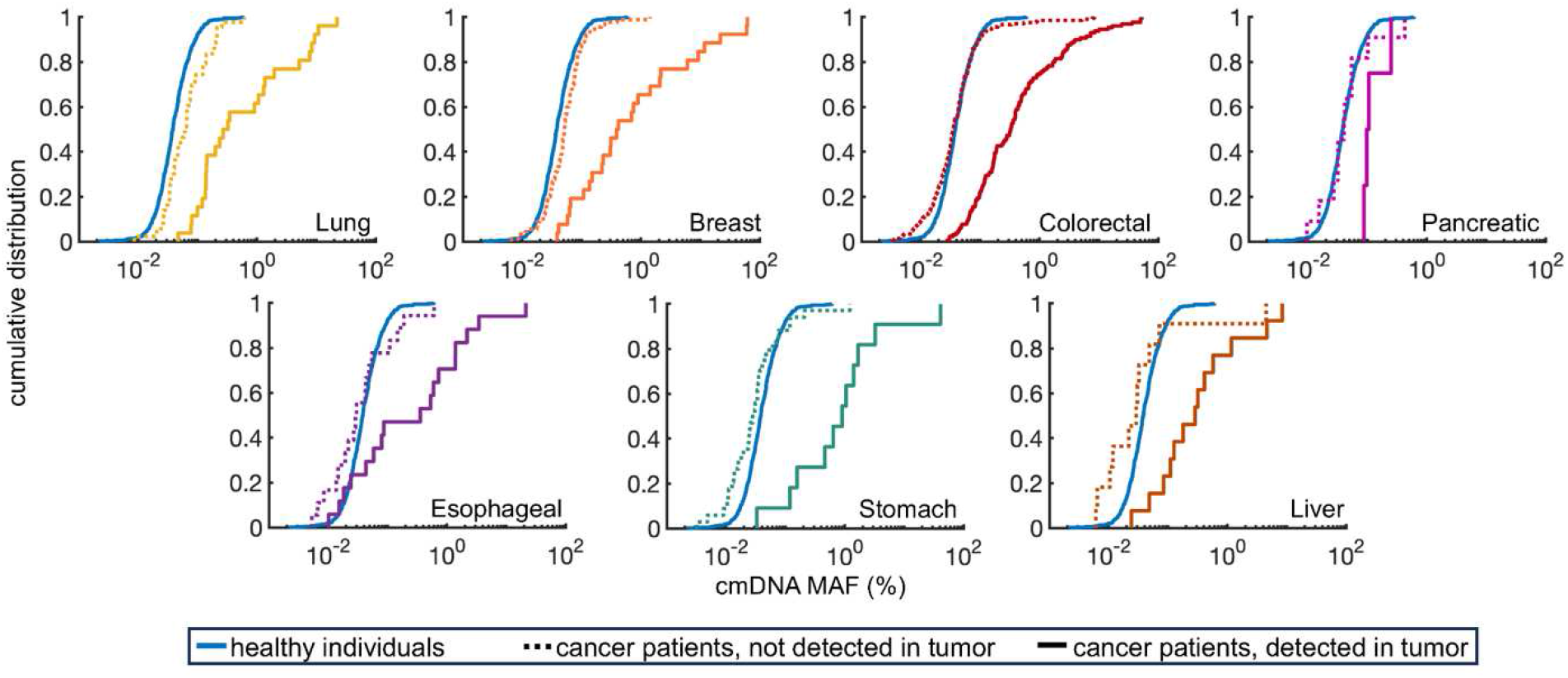
Cumulative distribution function of plasma mutant allele frequencies (MAFs) for the top plasma mutation of three groups: healthy individuals (solid blue line), patients of each cancer type whose top plasma mutation was not detected in the tumor (dotted line), and patients of each cancer type whose top plasma mutation was detected in the tumor (solid line). Sample sizes for each cancer type: lung (detected/detected: 43/26), breast (90/26), colorectal (205/129), pancreatic (11/4), esophageal (18/17), stomach (34/11), and liver (11/13). Ovarian cancer is not depicted since in the dataset of 7 patients, there was only one with the top mutation not detected in tumor.

**S10 Figure.**
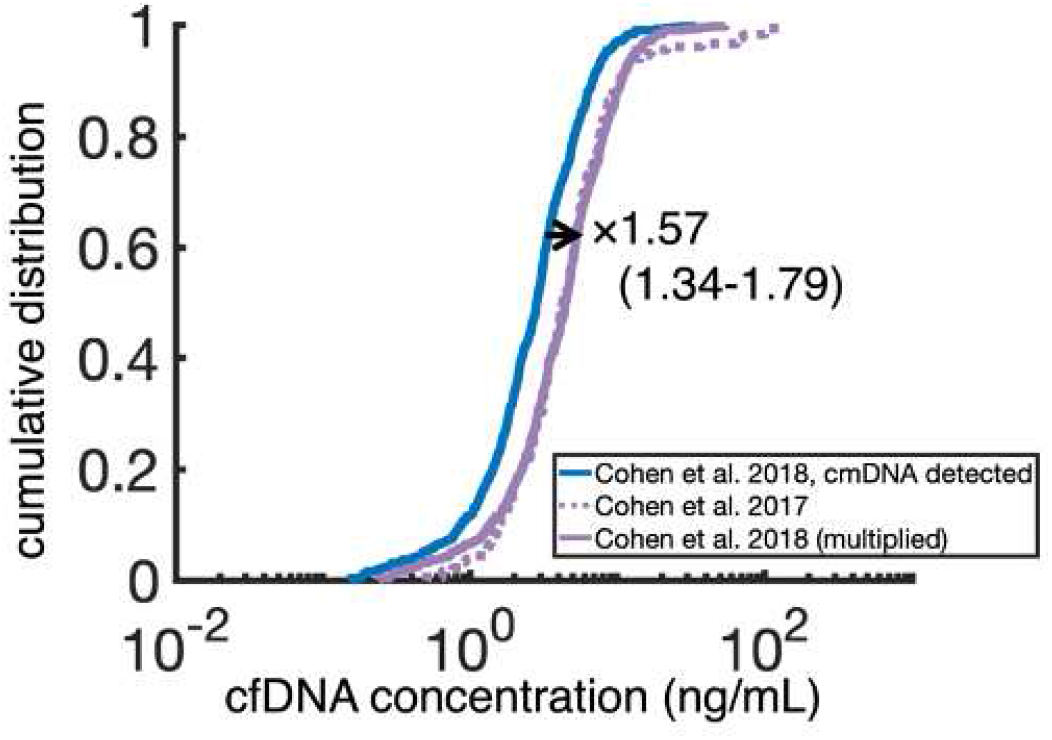
Multiplicative factor effect in cfDNA concentration between the 572 and 182 healthy individuals reported in the two studies by Cohen et al. of 2018 and 2017 respectively.

**S11 Figure.**
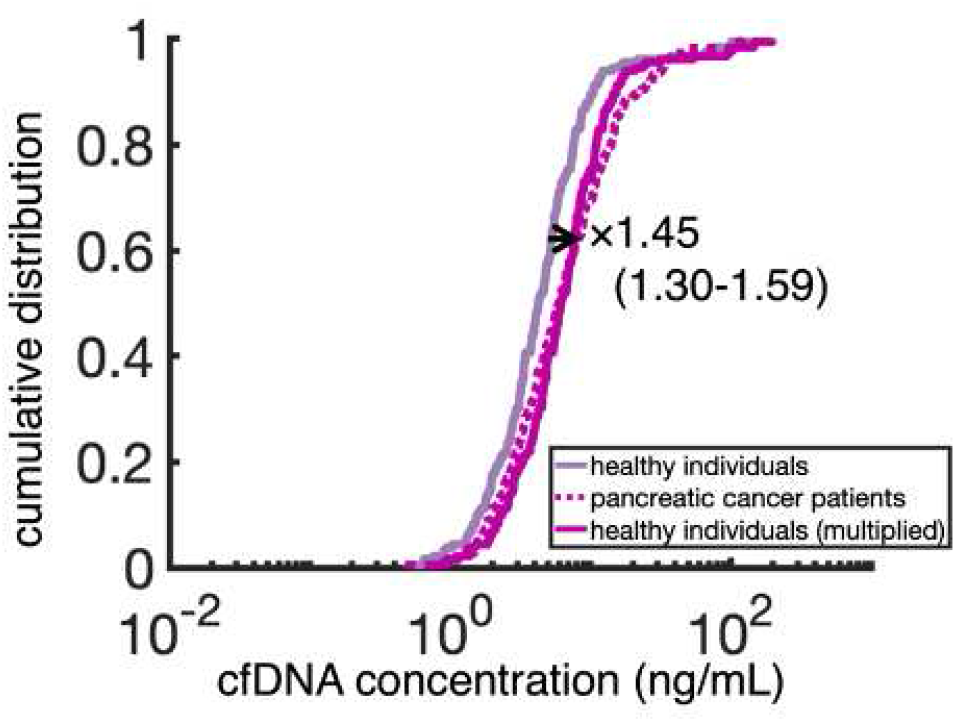
Multiplicative factor effect in cfDNA concentration between the 182 healthy individuals and the 221 pancreatic cancer patients reported in Cohen et al. 2017.

**S1a Table.** Cell-free DNA data from Cohen *et al*. 2018 [20] and Cohen *et al*. 2017 [40] analyzed in the manuscript.

| cfDNA dataset | data from [20] | exclude data included in [20] originating from [40] | exclude data with no cmDNA detected | data analyzed in Results | data from [40] analyzed separately in Discussion |
| --- | --- | --- | --- | --- | --- |
| healthy | 812 | 181 | 59 | 572 | 182 |
| lung cancer | 104 | - | 1 | 103 | - |
| breast cancer | 209 | - | - | 209 | - |
| colorectal cancer | 388 | - | 3 | 385 | - |
| pancreatic cancer | 93 | 46 | - | 47 | 221 |
| ovarian cancer | 54 | - | - | 54 | - |
| esophageal cancer | 45 | - | 1 | 44 | - |
| stomach cancer | 68 | - | 2 | 66 | - |
| liver cancer | 44 | - | - | 44 | - |

**S1b Table.** Summary of cfDNA dataset analyzed in Results, including sample size, age distribution (median and range), sex distribution (M/F), and cancer stage proportions (I/II/III) for each cancer type and healthy individuals.

**S1a Table.** Cell-free DNA data from Cohen *et al.* 2018 [20] and Cohen *et al.* 2017 [40] analyzed in the manuscript.
|  | cfDNA dataset<br>analyzed in<br>Results | Age in yrs<br>median (range) | Sex<br>M / F | Cancer Stage<br>I / II / III |
| --- | --- | --- | --- | --- |
| healthy | 572 | 36.5 (17–87) | 317 / 255 | - |
| lung cancer | 103 | 68 (43–86) | 62 / 41 | 46 / 27 / 30 |
| breast cancer | 209 | 57 (29–89) | 5 / 204 | 32 / 114 / 63 |
| colorectal cancer | 385 | 65 (22–93) | 210 / 175 | 76 / 191 / 118 |
| pancreatic cancer | 47 | 67 (44–86) | 28 / 19 | 2 / 39 / 6 |
| ovarian cancer | 54 | 60 (37–81) | 0 / 54 | 9 / 4 / 41 |
| esophageal cancer | 44 | 60 (30–82) | 39 / 5 | 5 / 28 / 11 |
| stomach cancer | 66 | 65 (35–82) | 51 / 15 | 20 / 29 / 17 |
| liver cancer | 44 | 62 (28–79) | 34 / 10 | 5 / 19 / 20 |

**S1c Table.** Summary of the cmDNA dataset. For each cancer type: the first column contains the number of patients whose mutations in tumor were reported; the remaining columns all pertain to these patients. The second column reports the number of patients with no plasma mutations detected (these patients are excluded from analysis in the Results). The last two columns report the number of patients for whom the top plasma mutation was not detected in the tumor versus also detected in the tumor.

|  | Patients with mutations in tumor reported | Patients with no mutations detected in plasma | Top mutation in plasma not detected in tumor | Top mutation in plasma also detected in tumor |
| --- | --- | --- | --- | --- |
| lung cancer | 70 | 1 | 43 | 26 |
| breast cancer | 116 | - | 90 | 26 |
| colorectal cancer | 336 | 2 | 205 | 129 |
| pancreatic cancer | 15 | - | 11 | 4 |
| ovarian cancer | 7 | - | 1 | 6 |
| esophageal cancer | 36 | 1 | 18 | 17 |
| stomach cancer | 47 | 2 | 34 | 11 |
| liver cancer | 24 | - | 11 | 13 |

**S2 Table.**
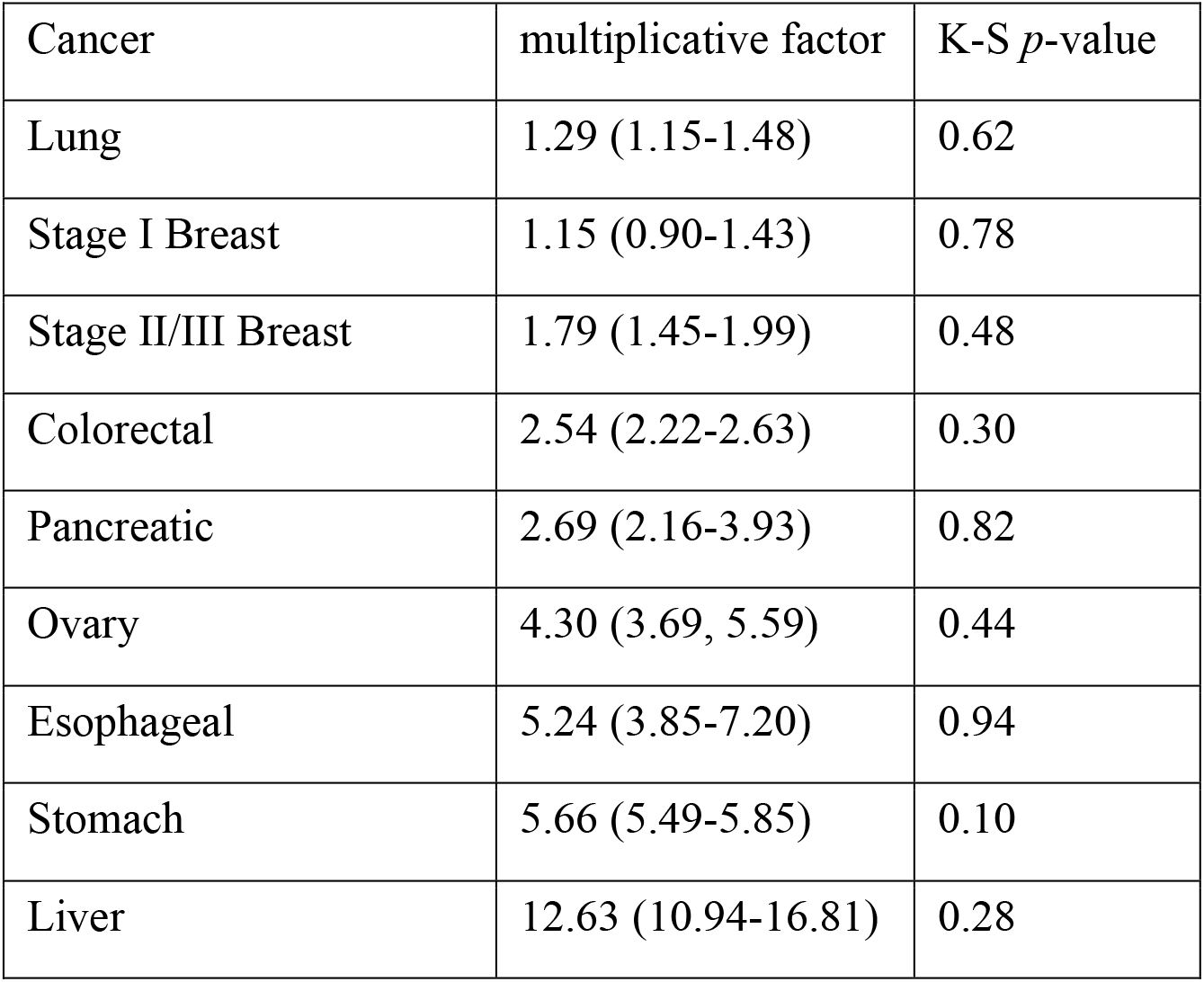
Multiplicative factors for cfDNA concentration, for each cancer type.

**S3 Table.** Multiplicative factors for cmDNA concentration not shed from tumor, for each cancer type.

| Cancer | multiplicative<br>factor | K-S<br>*p*-value | multiplicative<br>factor<br>MAF >0.01% | K-S<br>*p*-value | multiplicative<br>factor<br>MAF >0.015% | K-S<br>*p*-value | multiplicative<br>factor<br>MAF >0.02% | K-S<br>*p*-value |
| --- | --- | --- | --- | --- | --- | --- | --- | --- |
| Lung | 1.92<br>(1.42-2.52) | 0.89 | 1.93<br>(1.45-2.52) | 0.88 | 1.89<br>(1.41-2.45) | 0.88 | 1.75<br>(1.34-2.34) | 0.89 |
| Breast | 2.42<br>(2.12-2.76) | 0.41 | 2.50<br>(2.22-2.79) | 0.27 | 2.44<br>(2.16-2.66) | 0.31 | 2.35<br>(2.06-2.54) | 0.31 |
| Colorectal | 2.36<br>(1.92-2.66) | 0.51 | 2.49<br>(1.98-2.70) | 0.42 | 2.51<br>(2.00-2.74) | 0.45 | 2.59<br>(2.02-2.89) | 0.44 |
| Pancreatic | 4.01<br>(1.55-7.34) | 0.86 | 4.01<br>(1.43-8.53) | 0.82 | 4.01<br>(1.40-8.47) | 0.82 | 4.72<br>(1.39-9.35) | 0.78 |
| Esophageal | 3.31<br>(1.76-5.38) | 0.99 | 3.36<br>(1.77-6.07) | 0.99 | 3.64<br>(1.74-6.57) | 0.99 | 3.47<br>(1.52-6.78) | 0.99 |
| Stomach | 4.38<br>(2.74-5.77) | 0.86 | 4.56<br>(2.99-6.25) | 0.87 | 5.19<br>(2.98-7.70) | 0.83 | 5.01<br>(2.70-6.71) | 0.62 |
| Liver | 12.37<br>(5.54-24.62) | 0.99 | 14.56<br>(6.88-37.14) | 0.99 | 17.78<br>(7.19-46.06) | 0.99 | 16.96<br>(7.03-43.43) | 0.99 |

**S4 Table:** ROC-based performance metrics for cfDNA-driven cancer detection across cancer types and stages. The table reports AUC values with 95% confidence intervals, optimal cutoff points, sensitivity and specificity at the cutoff (with 95% CIs), and predictive values (PPV and NPV) at the cutoff (with 95% CIs).

| **Cancer Type** | **AUC**<br>**(95% CI)** | **Optimal**<br>**Cutoff** | **Sensitivity at**<br>**Cutoff**<br>**(95% CI)** | **Specificity at**<br>**Cutoff**<br>**(95% CI)** | **PPV**<br>**at Cutoff**<br>**(95% CI)** | **NPV**<br>**at Cutoff**<br>**(95% CI)** |
| --- | --- | --- | --- | --- | --- | --- |
| Lung | 0.608 (0.594,<br>0.622) | 2.457 | 0.763<br>(0.589, 0.778) | 0.422<br>(0.408, 0.485) | 0.179 (0.157,<br>0.200) | 0.890 (0.862,<br>0.916) |
| Lung, stage I | 0.616 (0.606,<br>0.633) | 2.633 | 0.758<br>(0.535, 0.838) | 0.452<br>(0.439, 0.526) | 0.091 (0.081,<br>0.105) | 0.949 (0.934,<br>0.965) |
| Breast, stage I | 0.556 (0.524,<br>0.575) | 2.455 | 0.813<br>(0.450, 0.833) | 0.377<br>(0.362, 0.518) | 0.060 (0.051,<br>0.068) | 0.965 (0.945,<br>0.983) |
| Breast, stages I, II | 0.710 (0.697,<br>0.723) | 3.697 | 0.655<br>(0.575, 0.727) | 0.668<br>(0.630, 0.705) | 0.368 (0.320,<br>0.412) | 0.860 (0.825,<br>0.893) |
| Colorectal | 0.784 (0.778,<br>0.788) | 4.105 | 0.742<br>(0.700, 0.766) | 0.704<br>(0.676, 0.718) | 0.629 (0.596,<br>0.669) | 0.799 (0.775,<br>0.820) |
| Colorectal, stage I | 0.809 (0.787,<br>0.828) | 4.081 | 0.809<br>(0.657, 0.861) | 0.693<br>(0.657, 0.739) | 0.269 (0.222,<br>0.318) | 0.960 (0.937,<br>0.980) |
| Pancreatic | 0.830 (0.814,<br>0.851) | 6.307 | 0.595<br>(0.420, 0.667) | 0.899<br>(0.738, 0.902) | 0.233 (0.158,<br>0.316) | 0.961 (0.945,<br>0.977) |
| Ovarian | 0.901 (0.890,<br>0.908) | 5.453 | 0.848<br>(0.670, 0.880) | 0.819<br>(0.797, 0.869) | 0.314 (0.267,<br>0.360) | 0.980 (0.967,<br>0.990) |
| Ovarian, stage I | 0.976 (0.968,<br>0.989) | 6.776 | 1.000<br>(0.608, 1.000) | 0.874<br>(0.862, 0.925) | 0.099 (0.025,<br>0.174) | 0.998 (0.994,<br>1.000) |
| Esophageal | 0.941 (0.936,<br>0.950) | 7.026 | 0.863<br>(0.748, 0.950) | 0.899<br>(0.888, 0.932) | 0.429 (0.373,<br>0.479) | 0.987 (0.981,<br>0.993) |
| Stomach | 0.918 (0.909,<br>0.928) | 5.111 | 0.879<br>(0.795, 0.940) | 0.794<br>(0.776, 0.834) | 0.340 (0.317,<br>0.361) | 0.981 (0.975,<br>0.989) |
| Stomach, stage I | 0.896 (0.883,<br>0.930) | 5.859 | 0.750<br>(0.550, 0.850) | 0.874<br>(0.759, 0.872) | 0.098 (0.054,<br>0.141) | 0.985 (0.976,<br>0.995) |
| Liver | 0.951 (0.944,<br>0.959) | 11.519 | 0.818<br>(0.688, 0.885) | 0.975<br>(0.968, 0.988) | 0.791 (0.670,<br>0.893) | 0.984 (0.977,<br>0.991) |

